# Major cell type differences between larval and adult hemichordate body plans

**DOI:** 10.1101/2025.10.31.685866

**Authors:** Paul Bump, Carolyn Brewster, Laurent Formery, Lauren Lubeck, Catherine Campbell, Maurizio Morri, Rene Sit, Daniel S. Rokhsar, Blair Benham-Pyle, Alejandro Sánchez Alvarado, Christopher J. Lowe

## Abstract

A major gap in our understanding of animal development is how adult body plans arise in animals with indirect development, where adults emerge from the transformation of a distinct larval form during metamorphosis. We address this question by examining cellular changes in the enteropneust hemichordate *Schizocardium californicum*, a species with a complex lifecycle and dramatic metamorphosis. Employing whole-body single-cell RNA sequencing, we chart the cellular composition and transcriptional dynamics of larval, metamorphosis, and adult stages. Our tissue level atlas reveals that ectodermal and endodermal cell types in larvae and adults occupy distinct transcriptional spaces, showing greater similarity to other cell types within the same life stage than to their counterparts in the opposite stage. In contrast, mesodermal cell types from both larvae and adults cluster closely together, indicating conserved transcriptional profiles. These findings demonstrate that the extensive morphological reorganization during metamorphosis is paralleled by profound shifts in cell-type specific transcriptional programs, highlighting the complexity of the larva-to-adult transition.

## Introduction

Our understanding of adult development is largely driven by the detailed study of a few select species, most of which are direct developers, defined by the formation of the adult body plan during embryogenesis (Held, 2005; McEdward & Janies, 1993; Raff, 2008). However, this type of developmental strategy does not represent the full diversity of animal life histories, and in many species the adult is formed through metamorphosis – the transformation of a larval body plan into an adult – often weeks to months after embryogenesis (Figure 1A) (Bolker, 1995; Hickman, 1999). Although species are largely categorized as either direct or indirect developers, developmental strategy is far less categorical and instead developmental mode largely operates along a continuum (Formery & Lowe, 2023; Wray, 2000). At one extreme are direct developers where adults form straight from the embryo and there are no intervening larval structures (Jäegersten, 1972; Nielsen, 1998; Pechenik, 1999). At the other end of the spectrum are species where the larval and adult body plans are developmentally and morphologically uncoupled with the adult formed by transformation of larval tissue during metamorphosis (McEdward & Janies, 1993). Most developmental model species lie closer to the direct end of the spectrum; even species with an ecologically significant larvae, like the frog *Xenopus laevis*, form the main elements of the chordate adult body plan during embryogenesis. On the other end of the continuum, sea urchins represent the extreme limit of indirect development: their embryos give rise to a bilaterally symmetric, planktonic larva that is both morphologically and axially very distinct from the radial adult body plan that forms by a drastic body reorganization during metamorphosis (Hyman, 1955; Wray & Raff, 1990).

**Figure 1.**
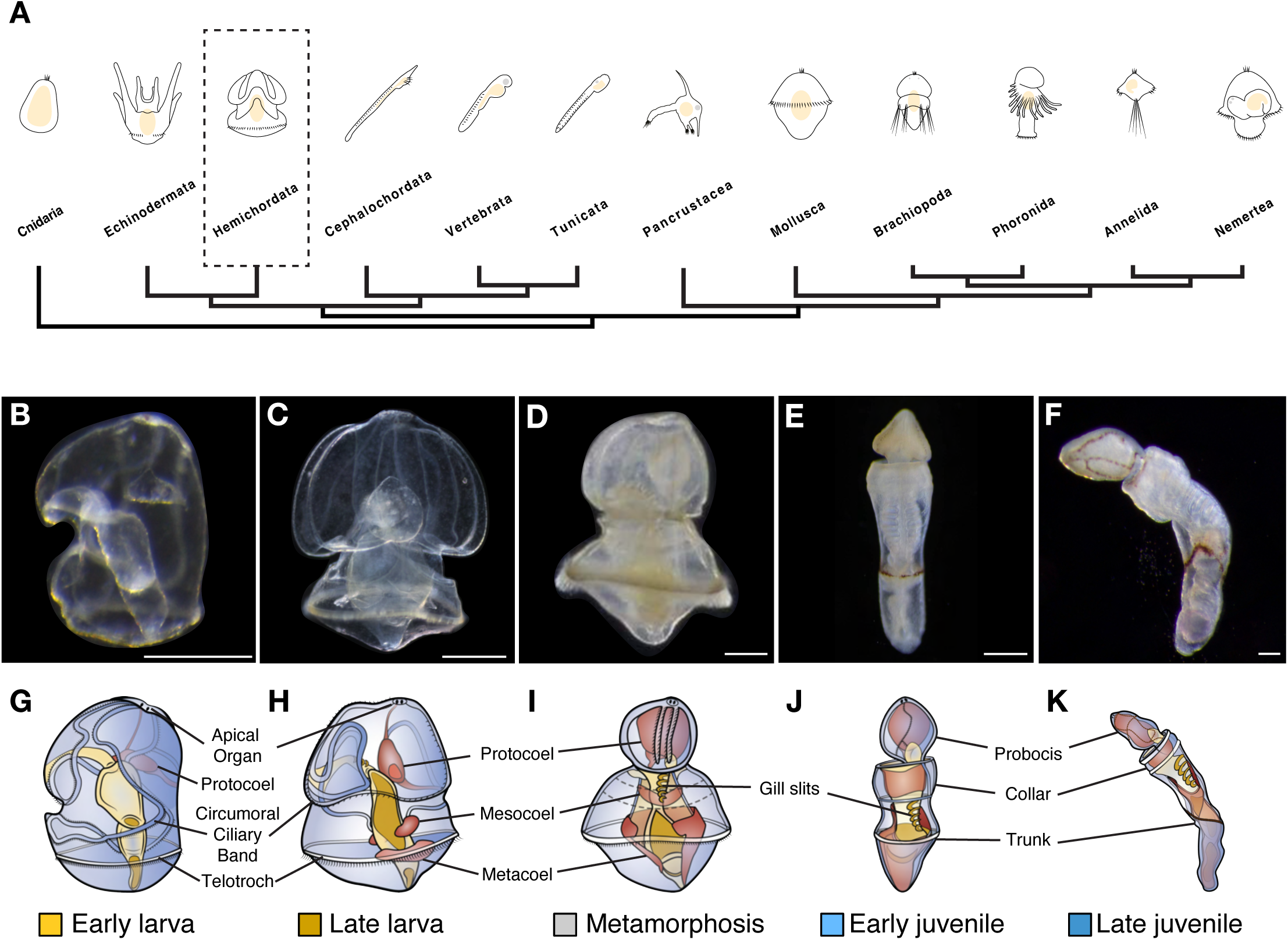
Indirect developing hemichordates as a model to study the transition between larval and adult body plans. A) Phylogenetic distribution of bilaterians highlighting the prevalence of larval types (Adapted from Formery and Lowe 2023). Light microscopy of the major life history stages of *S. californicum*: B) early larva, C) late larva, D) during metamorphosis, E) early juvenile, F) late juvenile. Schematic diagrams of the major life history stages of *S. californicum*: G) early larva, H) late larva, I) metamorphosis, J) early juvenile, K) late juvenile. In the schematics, ectoderm derivatives are shown in blue, endoderm derivatives in yellow, and mesoderm derivatives in red.

Indirect development has a broad phylogenetic distribution across bilaterians and has been proposed to represent the ancestral bilaterian developmental strategy (Martín-Zamora et al., 2023; Nielsen, 1998). However, our mechanistic understanding of animals defined by indirect development is largely limited to embryogenesis and early larval patterning. How the adult body plan forms by the transformation of the larval body plan rather than from embryonic development is poorly characterized (Formery & Lowe, 2023; Wray, 2000).

Recently, single-cell sequencing approaches have expanded our understanding of developmental trajectories and cell type diversification. However, the dynamics of cellular changes during metamorphosis remain poorly understood. Comparative work in single-cell genomics has either focused on direct developers or on embryonic and early larval stages, excluding the later adult life history stages in animals with complex life cycles (Foster et al., 2022; Massri et al., 2021; Paganos et al., 2021; Perillo et al., 2020; Piovani et al., 2023). For a comprehensive understanding of developmental mechanisms through metamorphosis, it is essential to have insights into the cellular composition of larvae and adults and how these may change through metamorphosis. Recent studies are beginning to reveal that this perspective is essential to fully evaluate the cellular diversity represented by particular lineages, to test hypotheses of cell type evolution, and to understand the evolution of animal body plans (Link et al., 2025; Paganos et al., 2025; Sebé-Pedrós, Chomsky, et al., 2018; Sebé-Pedrós, Saudemont, et al., 2018).

The enteropneust hemichordate *Schizocardium californicum* is an excellent model for the study of indirect development (Cameron & Perez, 2012) (Figure 1B-F). Like many enteropneusts, *S. californicum* develops via a tornaria larva that is organizationally similar to echinoderm larvae, with two primary ciliary bands used for swimming and feeding, and limited musculature (M. Hadfield & Strathmann, 1996; Lacalli & Gilmour, 2002; Morgan, 1891). This type of larva has been termed “head larvae” as their body plan is largely defined by regulatory networks that define an anterior “head” territory in bilaterians and are missing a Hox-patterned trunk domain (Gonzalez et al., 2016; Lacalli, 2005; R. R. Strathmann, 2020). The formation of the adult through a radical metamorphosis transforms the planktonic larva into a burrowing benthic adult and involves extensive remodeling of the ectoderm, rapid expansion of the mesoderm, reorganization of the endoderm, and the formation and extension of the trunk territory. Adults are adapted for a burrowing lifestyle using a muscular proboscis, and feed using both mucociliary particle capture and filter feeding through their pharyngeal gill slits (Cameron, 2002; Gonzalez & Cameron, 2009). The tripartite gut is structurally similar between larva and adult, but the gill slits begin to develop in the late larva in the anterior gut and perforate the anterior trunk of the adult after a posterior migration during metamorphosis (Nielsen & Hay-Schmidt, 2007). The larval nervous system is defined by an apical organ and neurons associated with the feeding ciliary band, whereas the adult has an extensive basiepidermal plexus and both a dorsal and ventral nerve cord (Andrade López et al., 2023; Bullock, 1945; Kaul-Strehlow et al., 2015; Miyamoto et al., 2010).

While morphological changes during enteropneust metamorphosis are extensive, the axial register between life histories is maintained along with the original mouth and anus. Previous work suggests that metamorphosis does not involve extensive larval cell death and larval cells may be carried over into the adult (Bump et al., 2022), raising the question of the fate of larval cells in the adult body plan.

The goal of this study is to identify similarities and differences in cellular composition of the larval and adult body plan of *S. californicum* using single-cell transcriptomics at time points representing larval, metamorphosis, and juvenile stages. We assess the degree to which cell types overlap between life histories and whether metamorphosis represents not only a major morphological change in body plan organization, but also a change in cellular composition. We find that larval and juvenile cells largely occupy distinct transcriptional states and show greater similarity to other tissues within the same life-history stage than to their putative counterparts across stages. Additionally, pulse-chase EdU experiments through metamorphosis demonstrate persistence of labeled larval cells into juveniles, arguing against wholesale larval cell replacement and supporting cellular reprogramming during metamorphosis.

## Results

### Sampling distinct life history stages in the same organism

To understand the cell types that make up distinct larval and juvenile animals, we sampled five distinct life history stages in *S. californicum*: early larva, late larva, metamorphosis, early juvenile, and late juvenile based on the previous description of development (Gonzalez et al., 2018) (Figure 1B-F) and which are generally representative of other, more distantly related indirect developing enteropneust species such as *Ptychodera flava* and *Balanoglossus simodensis*.

#### Early larva

The first larval stage was selected at 22 days post fertilization (Figure 1B, 1G). The larval body plan is fully formed but lacks any nascent adult structures. Tornaria larvae are small, simple planktonic organisms that feed and swim using cilia organized into two distinct ciliary bands (Figure 1B, 1G): the circumoral ciliary loop lines the anterior of the larva and is used primarily for feeding, while the circumferential telotroch (Figure 1G, 1H) in the posterior of the larva is defined by long compound cilia and is exclusively used for locomotion (Garstang, 1939; Lacalli & Gilmour, 2002; R. Strathmann & Bonar, 1976). The remaining ectoderm is squamous epithelium with a ciliated neurotroch that runs along the ventral midline between the anus and the posterior circumoral ciliary loop. At the most dorsal anterior there is an apical tuft of cilia and an associated underlying apical organ with neurons that project throughout the epithelium and two pigmented eyespots (Gonzalez et al., 2018). There is very little mesoderm in early larvae, as only a single anterior protocoel is present at this stage (Figure 1G). The larval muscle is localized around the pharynx and in a single apical, muscular strand that contracts and connects the protocoel to the apical plate (R. Strathmann & Bonar, 1976; Tagawa et al., 1998) (R. Strathmann & Bonar, 1976; Tagawa et al., 1998). The endoderm of the tornaria larva consists of a tripartite gut composed of the pharynx, stomach, and intestine with a terminal anus.

#### Late larva

The second larval stage was selected after two months of growth (Figure 1C, 1H), when the larva is competent to begin metamorphosis (M. G. Hadfield et al., 2001). While morphologically similar to the early tornaria, this stage is defined by a much larger size and more extensive feeding ciliary band and evidence of pre-patterning of adult structures. In the mesoderm the additional two pairs of posterior coeloms, the mesocoels and metacoels (Figure 1H), have formed, and nascent gill slits are present in the anterior endoderm. The posterior to the telotroch is larger and there is an expanded posterior region or anal field. The nervous system is extensive, with an expansion of the serotonergic nervous system throughout the ectoderm and the first signs of the development of the adult dorsal cord (Gonzalez et al., 2018).

#### Metamorphosis (Agassiz stage)

The third developmental stage selected was the midpoint of metamorphosis (Figure 1D, 1I). Metamorphosis is an approximately 48 hour long process that begins with a thickening of the larval ectoderm, the initiation of trunk development, and an expansion of all the mesodermal coeloms as the large blastocoelar space is reduced in an anterior to posterior wave (Agassiz, 1873; Nielsen & Hay-Schmidt, 2007). The midpoint of metamorphosis contains morphological elements of both the larval and adult body plans. It is defined by a thickened prospective proboscis ectoderm resulting from the degeneration of the circumoral ciliary band, and an expanded anterior coelom that becomes the musculature of the proboscis. The stomochord has formed as an anterior projection from the gut at the base of the proboscis supporting the heart/kidney complex responsible for coelomic fluid filtration and circulation (Balser & Ruppert, 1990). At this stage trunk formation has been initiated and elongation has begun, but the blastocoel remains prominent even as the mesocoel and metacoels expand (Figure 1D, 1I). In the endoderm the nascent gill slits are migrating posteriorly towards the trunk and the collagenous skeletons of the proboscis and gill slits are beginning to develop. Metamorphosing *S. californicum* are still planktonic and use the telotroch for locomotion.

#### Early and late juveniles

The definitive adult body plan is represented in our study by early (Figure 1E, 1J) and late (Figure 1F, 1K) juveniles: benthic, burrowing worms that have not reached sexual maturity. The tripartite adult body plan typical of enteropneusts is clearly defined into proboscis, collar, and trunk regions: the mouth opens ventrally at the base of the proboscis, the gill pores perforate the ectoderm in the anterior trunk, and both dorsal and ventral nerve cords are present. Early juveniles are newly settled animals with well-developed proboscis and collar that have not undergone substantial trunk elongation (Figure 1E, 1J). To ensure that we captured the full cellular complement of the adult body plan, we also sampled late juveniles (Figure 1F, 1K) defined by a significant elongation of the trunk.

### Single-cell sequencing reveals the diversity of hemichordate cell types

To understand the cellular composition of each of these distinct life history stages whole organism cell suspensions were prepared for single-cell RNA sequencing (scRNA-seq). Cell suspensions were processed and a total of 87,021 single-cell profiles were sequenced: 15,403 cells for early larva, 22,027 for late larva, 13,771 during metamorphosis, 20,061 for early juvenile, and 15,759 for late juvenile.

We generated an atlas of *S. californicum* across life history stages by clustering cellular profiles, identifying 56 clusters containing between 68 and 8,392 cells (average 1,554 cells/cluster) (Figure 2A, Supplemental Figure 1A, Supplemental Table 1). Cell clusters are defined by markers obtained by analysis of differential expression (log fold change and p-value) and specificity (Supplemental Table 2). Based on cluster-specific gene expression we further organized these cell type clusters into 12 cell classes (cartilage, ciliary band, endoderm, epidermis, immune cells, mesoderm, muscle, neural, pigment, proliferative, secretory, and unknown) (Figure 2B, Supplemental Table 2).

**Figure 2.**
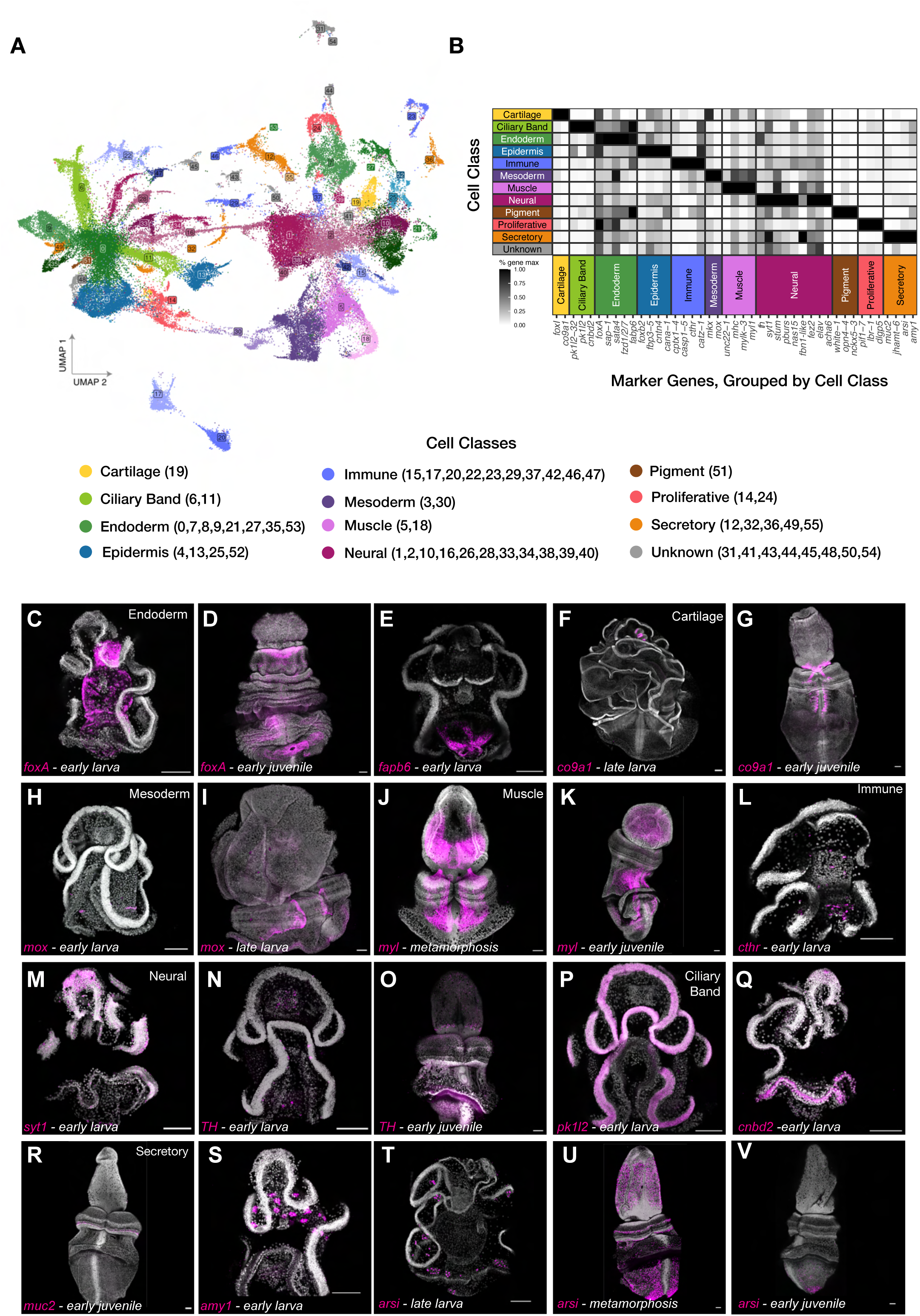
Single-cell sequencing reveals the broad diversity of hemichordate cell types. A) UMAP embedding of the cell classes of *S. californicum;* 55 clusters were identified and 12 cell classes were described. B) Markers of cell classes of *S. californicum*, percentage of gene max was calculated by taking the mean normalized expression of the gene of interest and dividing by the max for any cell class. C-T) HCR signal in magenta, nuclei are in grey, scale bars are 100 µm. C) *foxA* HCR expression in early larva. D) *foxA* HCR expression in early juvenile. E) *fabp6* HCR expression in early larva. F) *co9a1* HCR expression in late larva. G) *co9a1* HCR expression in early juvenile. H) *mox* HCR expression in early larva. I) *mox* HCR expression in late larvae. J) *myl* HCR expression during metamorphosis. I) *myl* HCR expression in early juvenile. L) *cthr* expression in early larva. M) *syt1* HCR expression in early larva. N) *TH* HCR expression in early larva. O) *TH* HCR expression in early juvenile. P) *pk1l2* HCR expression in early larva. Q) *cnbd2* HCR expression in early larva. R) *muc2* HCR expression in early juvenile. S) *amy1* HCR expression in early larvae. T) *arsi* HCR expression in late larvae. U) *arsi* HCR expression in during metamorphosis. V) *arsi* HCR expression in early juvenile.

We validated our cell type clusters using both newly discovered markers from our cluster annotations (Supplemental Table 2, 3) and known tissue markers in hemichordates, by plotting the relative expression of a gene of interest in each cell class (Figure 2B) and then performing HCR (hybridization chain reaction) fluorescent *in situ* hybridization (Figures 2C-T, Supplemental Table 4).

#### Endoderm derivatives

*Endoderm clusters: 0, 7, 8, 9, 21, 27, 35, 53* (Figure 2C-G, Supplemental Figure 2A-C). Differentially expressed genes for endoderm include fatty acid binding proteins and *foxA*, a known endoderm marker in hemichordates (Fritzenwanker et al., 2014; Green et al., 2013). *FoxA* is found in all eight of our endoderm clusters, with limited expression outside of this group. It is expressed throughout the tripartite gut of *S. californicum* larvae in the mouth, stomach, and intestine (Figure 2C and 2D), consistent with prior observations in *S. kowalevskii* (Fritzenwanker et al., 2014). As in *S. kowalevskii, S. californicum* juveniles have a ring of *foxA* expression in the anterior collar groove, but unlike *S. kowalevskii* there also seems to be a broader expression domain in the ectoderm (Figure 2D). We also identified additional markers from our single-cell analysis that enabled finer-scale characterization of endoderm. For example, the gut marker *fabp6*, which encodes a fatty acid binding protein family member responsible for lipid balance and retinol- and retinoic acid-binding across animals (Y. Zheng et al., 2013), is highly expressed in the larval intestine (Figure 2E) while *fzd1/2/7* was expressed in the larval stomach (Supplemental Figure 2C).

*Cartilage cluster: 19* (Figure 2F-G, Supplemental Figure 2D-F). A single cluster was enriched for several cartilage markers, including *collagen alpha-1(IX) chain-like* (*co9a1*). Since cartilage in enteropneust hemichordates is secreted by the endodermal pharyngeal epithelium (Rychel et al., 2006) and the gill slits specifically are made up of fibrillar collagen (Pardos & Benito, 1988), we expected *co9a1* expression would be localized to the gill slits, a characteristic deuterostome feature. We found that *co9a1* is expressed in a small pocket of cells in the late larva in the developing gill slits (Figure 2F), through metamorphosis (Supplemental Figure 2F), and in juveniles in the gill bars and the wishbone-shaped proboscis skeleton (Figure 2G). This cluster also expresses *foxI,* a marker implicated in gill pouch endoderm and in the posterior gut (Supplemental Figure 2E) (Fritzenwanker et al., 2014).

#### Mesoderm derivatives

*Mesoderm clusters: 3 and 30* (Figure 2H-I, Supplemental Figure 3A-B). Differentially expressed genes in the mesoderm clusters included *mesenchyme homeobox* (*mox*), a gene expressed in the paired posterior coelomic cavities in *S. kowalevskii* (Lowe et al., 2006). Using HCR we detected *mox*-positive cells in *S. californicum* during initiation of the posterior coelom formation prior to any clear morphological outpocketing of mesoderm (Figure 2H) as well as in clearly defined coeloms of the late larva (Figure 2I).

*Muscle clusters: 5 and 18* (Figure 2J-K, Supplemental Figure 3C-E). Enteropneust adults are highly muscular, and muscle clusters were identified in our data set by the expression of muscle components such as *titin, popeye domain-containing protein 3,* and *myosin light chain (myl*) (Supplemental Figure 3D, E). Expression *myl* was detected in all life stages and was particularly prevalent near the end of metamorphosis where longitudinal fibers are forming to give rise to the musculature of the juvenile worm (Figure 2J, 2K).

*Immune clusters: 15, 17, 20, 22, 23, 29, 37, 42, 46, 47* (Figure 2L, Supplemental Figure 4A-B). scRNA-seq allowed us to characterize many previously undescribed cell types including putative immune cells. We found a number of immune populations that expressed macrophage mannose receptors including C-type lectins expressed by most tissue macrophages and dendritic cells (Taylor et al., 2005; Vasta et al., 2017), as well as a specific population of cells (cluster 17) marked by the expression of *collagen triple helix repeat-containing protein* (*cthr*) (Figure 2L), a secreted glycoprotein implicated in tissue repair processes (Myngbay et al., 2021).

#### Ectodermal derivatives

*Neural clusters: 1, 2, 10, 16, 26, 28, 33, 34, 38, 39, 40* (Figure 2M-O, Supplemental Figure 5A-C). We identified neural cell clusters by the expression of *synaptotagmin* (*syt1*), which is involved in synapse docking (Littleton et al., 1993) (Figure 2M) as well as *elav*, a neuron-specific RNA binding protein expressed in differentiated bilaterian neurons (Cunningham & Casey, 2014; Koushika et al., 1996; Robinow & White, 1991), but as noted in other species, it does not mark all neurons (Berger et al., 2007; Pham & Hobert, 2019). Neural subtypes were identified by the expression of enzymes involved in the synthesis of neurotransmitters or transporters: *tyrosine hydroxylase* (*TH*), which marks dopaminergic neurons (Figure 2N and 2O), the GABAergic marker *GAD*, and cholinergic marker *VAChT* (Supplemental Figure 5C). We observed these neural cell types in both larvae and juveniles.

*Ciliary band clusters: 6 and 11* (Figure 2P-Q, Supplemental Figure 5E-G). The tornaria larva is characterized by two distinctive ciliary bands, the circumoral ciliary band and the telotroch (Figure 1B, 1C, 1G, 1H). Clusters 6 and 11 expressed known ciliary markers. Cluster 6 showed enrichment of *polycystic kidney disease protein 1-like 2* (*pk1l2*), which has been associated with the primary apical cilium of renal epithelia (Marra et al., 2019; Pfeffer et al., 1998; Yoder et al., 2002). When we examined the expression of *pk1l2* via HCR imaging we found that it labeled the circumoral ciliary band but not the telotroch (Figure 2P). Cluster 11 was marked by *cyclic nucleotide-binding domain containing protein* (*cnbd2*), which may be involved in ciliary beating (Darszon et al., 2008; Hildebrandt et al., 1997). HCR imaging showed that *cnbd2* had a distinct expression pattern in the telotroch (Figure 2Q).

*Secretory clusters: 12, 32, 36, 49, 55* (Figure 2R-V, Supplemental Figure 6A-B). Hemichordate epidermis is highly secretory, producing large amounts of mucus from the epidermis, and we found several secretory clusters in both larvae and juveniles. We propose that cluster 12 is a secretory cell population based on the high expression of *mucin2 (muc2)* (Supplemental Figure 6B). Mucins have been implicated as important for muco-ciliary feeding in hemichordates (Ruppert et al., 1999). HCR imaging showed *muc2* expressed in a thin circumferential band of cells in the collar of the juvenile (Figure 2R), similar to the expression of mucins in *S. kowalevskii* (Simakov et al., 2015). Even for putative secretory clusters with a smaller number of cells (374 for cluster 49) we were able to recover distinct patterns of cell type-specific gene expression. Cluster 49 is characterized by expression of *alpha-amylase* (*amy1*). Alpha-amylases hydrolyze 1,4-alpha-glucoside bonds in oligosaccharides and polysaccharides in the salivary gland of humans (Samuelson et al., 1988). HCR imaging showed *amy1* expression in a very distinctive pattern of cells surrounding the larval stomach (Figure 2S), so we propose that this cell population may be involved in secretory function. Finally, we found a cluster of secretory cells with a striking shift in abundance and distribution over developmental time. The markers of a putative secretory cluster 32 included *arylsulfatase* (*arsi*), a protease that is upregulated in the *Xenopus laevis* tadpole tail as an integral part of the resorption process during metamorphosis (Das et al., 2006). In the late larva, these *arsi*+ cells are found in clusters throughout the ectoderm (Figure 2T), become broadly distributed during metamorphosis (Figure 2U), and finally are predominantly restricted to the most posterior of the early juvenile worm (Figure 2V). Future work will be necessary to investigate the function of this intriguing cell type during metamorphosis.

### Divergence of transcriptional states between life histories

Given that we had sampled the single-cell composition of five life history stages with distinct morphologies and ecological demands, we wanted to understand how cellular composition relates to life history. We considered three possible scenarios: First, that larval cell types represent a subset of adult cell types; second, that the cells of larval and adult cells share transcriptional identity, while each stage also possesses unique cell types; and third, that larval and adult body plans are composed of distinct cell types with limited overlap in transcriptional identity.

To test these possibilities, we plotted life history on our scRNA-seq clusters (Figure 3A). We observed a striking pattern when comparing the distribution of cells across the five different life history stages: larval and juvenile stages largely clustered in distinct regions of transcriptional space (Figure 3A). To confirm that this observation was not an artifact resulting from our chosen methods for integrating cells from different stages into a single transcriptional space (See Material and Methods, Supplemental Figure 1), we performed “Harmony” and “Portal” integrations, which still resulted in spatially separated clustering of larval and juvenile life stages (Supplemental Figure 1B and 1C).

**Figure 3.**
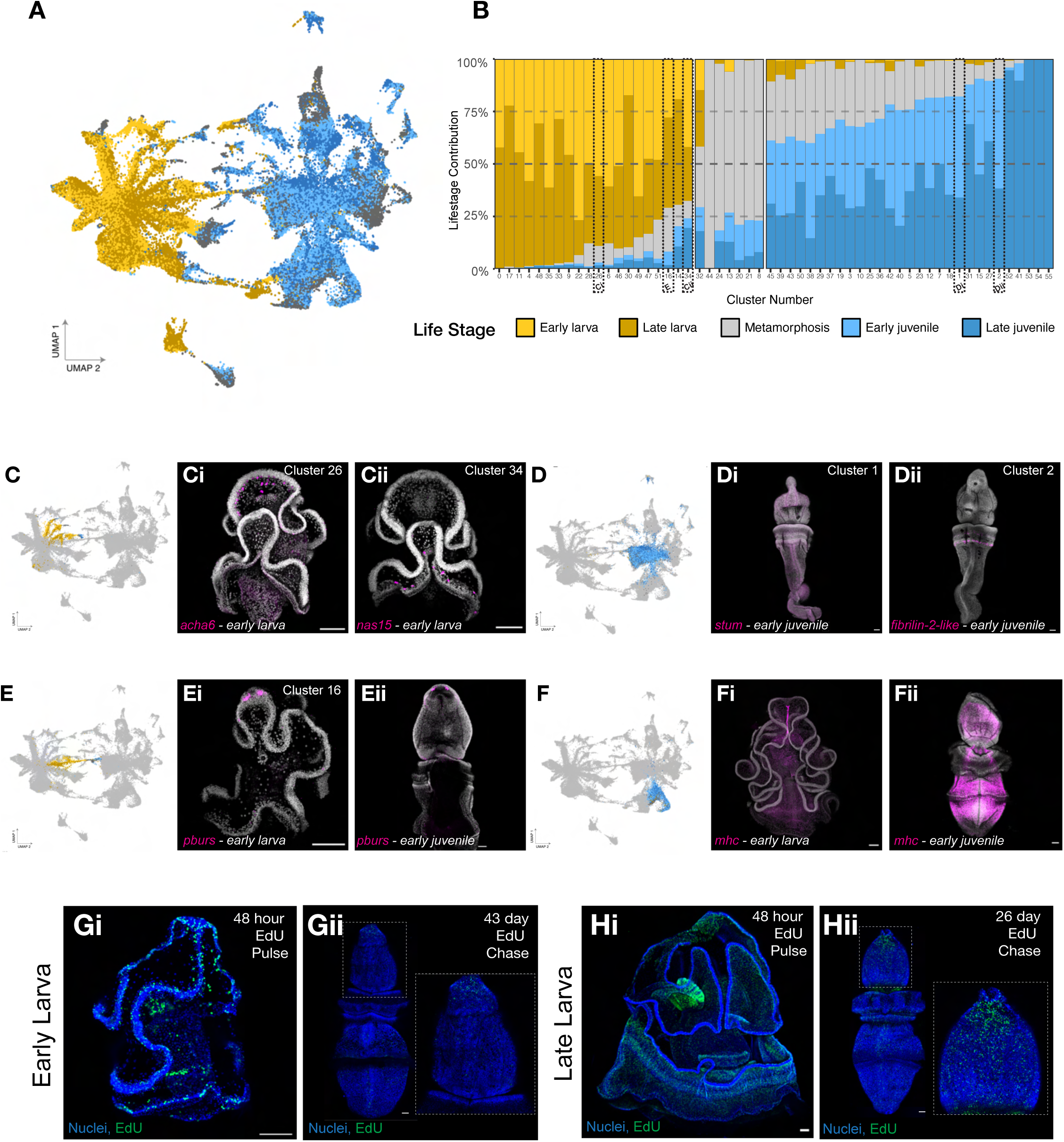
Divergence of transcriptional states is consistent with morphological changes between life history stages. A) UMAP embedding of *S. californicum* single-cell RNA-seq data with cells colored by life stage. Yellow: early larva, dark gold: late larva, grey: metamorphosis, light blue: early juvenile, dark blue: late juvenile. B) Bar chart of life stage composition of clusters. C) UMAP embedding with larval neural clusters (26, 28, 33, 34) colored by life stage. Ci) *acha6* HCR expression in early larvae. Cii) *nas15* HCR expression in early larvae. D) UMAP embedding with juvenile neural clusters (1, 2) colored by life stage. Di) *stum* HCR expression in early juvenile. Dii) *fibrilin-2-like* HCR expression in early juvenile. E) UMAP embedding with neural cluster 16 colored by life stage. Ei) *pburs* HCR expression in early larvae. Eii) *pburs* HCR expression in early juvenile. F) UMAP embedding with muscle clusters (5, 18) colored by life stage. Fi) *mhc* HCR expression in early larvae. Fii) *mhc* HCR expression in early juvenile. Gi) EdU pulse of early larva for 48 hours. Gii) EdU chase after 43 days once larva have completed metamorphosis, inset is of proboscis region of the juvenile worm, boxed. Hi) EdU pulse of late larva for 48 hours. Hii) EdU chase after 26 days once larva have completed metamorphosis, inset is of proboscis region of the juvenile worm, boxed.

To examine changes in cell type compositions between life stages, we looked at the proportion of cells in a given cluster from each timepoint (Figure 3B). Broadly, these life stage contributions matched our expectations of the morphology of the organism: some tissues are composed almost entirely of juvenile cells, like cartilage (cluster 19), while others are predominantly larval cells, such as the ciliary band (clusters 11 and 6). Consistent with the stage-related decrease in the number of cells in ciliary band clusters, we showed that the expression of markers of cluster 6 such as *pk1l2* decrease over time and are restricted to the circumoral ciliary band and future oral groove of the proboscis during metamorphosis (Supplemental Figure 5G). This aligns with known morphology and with our previous study that found a higher level of cell death in the circumoral ciliary band (Bump et al., 2022).

Based on our annotation, neural cells are one of most broadly distributed classes of cell types in both larvae and juveniles. Some cell types were present in both life stages but varied in their life stage composition, with a key example being the neural cell types (clusters 26, 16, 34, 1, 2 dashed boxes in Figure 3B with corresponding panels in Figure 3C-E).

There were neural clusters composed of predominately larval cells (Figure 3C), such as cluster 26 (Figure 3Ci) which was marked by expression of *nas15*, a zinc metalloproteinase, which is a part of a large family of astacins, some of which are expressed in the nervous system (Park et al., 2010). This cluster also expressed *lysine-specific demethylase 8* (*kdm8-7*), an important regulator of neurite morphogenesis (Zibetti et al., 2010). We found *nas15* expressed throughout the posterior of the larva in cells of the developing telotroch (Figure 3Ci). This neural cell type is absent in juvenile life stages, likely due to the loss of ciliary band at metamorphosis. Neural cluster 34 expressed *dopamine beta-hydroxylase* (*dopo*), which converts dopamine to noradrenaline (Axelrod, 1972) and *neuronal acetylcholine receptor subunit alpha 6* (*acha6*), which belongs to the ligand-gated ion channel superfamily, including GABA, glycine, and 5-HT3 receptors (Betz, 1990). We found *acha6+* cells distributed around the ventral anterior of the larva (Figure 3Cii). Other neural cell clusters were composed of cells that were predominantly from the juvenile stage, such as clusters 1 and 2 (Figure 3D). Cluster 1 was marked by the expression of *stum*, which has been implicated in mediating mechanical sensing in receptor neurons and is a transducer for mechanical stimuli (Desai et al., 2014). HCR showed *stum* expression along the dorsal and ventral nerve cords of the juvenile and projecting into the proboscis on the dorsal surface (Figure 3Di). Neural cluster 2 is marked by *fibrilin-2-like*, a large modular structural glycoprotein component of the extracellular matrix of peripheral nerves (Chernousov et al., 2010). Visualizing *fibrilin-2-like* via HCR showed expression in the collar of juveniles, with an enrichment of *fibrilin-2-like*+ cells with neural morphology at the base of the collar (Figure 3Dii). An example of a neural cell type present in both life history stages, neural cluster 16 (Figure 3E) is marked by *partner of bursicon* (*pburs*). HCR of *pburs* revealed expression in the eyespots as part of the larval apical organ (Figure 3Ei) (Nakajima et al., 2004; Nielsen, 2005). As development continued, expression was persistent in the eyespots (Supplemental Figure 5D) and continued into the juvenile in a persistent expression territory in two patches at the tip of the proboscis (Figure 3Eii). While the apical tuft of cilia associated with the apical organ degenerates at metamorphosis, some of the neurons and the eyespots appear to persist in the juvenile body, at least at juvenile stages (Gonzalez et al., 2018).

Finally, as described above, while many cell types had larval- or juvenile-specific transcriptional signatures, muscle (clusters 5 and 18) was an interesting exception (Figure 3F). A highly differentially expressed gene in these muscle clusters is *myosin heavy chain* (*mhc*). One of the few muscle types that is present in the larva is the apical strand muscle, which is marked by the expression of *mhc* (Figure 3Fi). During metamorphosis and into the early juvenile stage, *mhc* expression expands into the distinctive musculature of the juvenile worm, one of its defining morphological characters (Figure 3Fii, Supplemental Figure 3F). While the prevalence and distribution of muscle cells change over developmental time in the animal, our single-cell analysis suggests that transcriptional signature of these clusters remains essentially unchanged as they do not show spatially separated clustering of larval and juvenile cells (Figure 3F).

Given the changes that we saw in the transcriptional profile, we considered two principal possibilities: (1) extensive larval cell death during metamorphosis followed by proliferation of adult stem cell populations, or (2) larval cells are carried over into the juvenile animal with modified patterns of gene expression. Extensive larval cell death at metamorphosis is not supported by our previous observations (Bump et al., 2022), so to further explore the possibility of cells being carried over from larval to juvenile we performed a series of EdU (5-ethynyl-2’-deoxyuridine) pulse-chase experiments through metamorphosis. We incubated early and late larvae with EdU for 48 hours to label any cell in S-phase (Figure 3G, 3H), then washed out the EdU label and let the animals continue to grow and develop until they completed metamorphosis. In an early larva, a 48-hour pulse indiscriminately labels a number of different cellular populations including those in the ectoderm (ciliary band), mesoderm (wrapped around the gut), and endoderm (in the mouth and gut) similar to what we had observed with a short EdU pulse previously (Bump et al., 2022). Forty-three days later, we found that some EdU+ cells were still present, particularly in the anterior proboscis and along the dorsal midline (Figure 3Gii), indicating that at least some cells from the larval body plan persist through to the juvenile stage. In late larvae, closer to metamorphosis, 48-hour pulses label many cell populations within all three germ layers (Figure 3Hi). These cell population include the ciliary bands, the epithelium of the presumptive collar and proboscis ectoderm, the mesocoel and metacoel, and a large enrichment in the gill bars, consistent with previous reports of shorter EdU pulse-chase experiments at this particular stage (Bump et al., 2022). In this case, EdU+ cells persisted in the juvenile, with a strong enrichment in proboscis, collar, and dorsal cord (Figure 3Hii). From this we conclude that larval cells could be carried into the juvenile animal with possible changes in transcriptional profile.

### Differences between cells in larval and adult body plans

We further quantified and interrogated the overall transcriptional relationships between cell types and life stages using our scRNA-seq dataset. While two- or three-dimensional embeddings like UMAP plots (Figure 2A, Figure 3A) are convenient for visual analysis, low-dimensional representations unavoidably distort complex relationships among large numbers of clusters (Healy & McInnes, 2024). To more accurately quantify similarity (closeness) or dissimilarity (remoteness) of clusters in 55-dimensional principal component (PC) space, we calculated the centroid for the PC embeddings of the cells in each cluster and then calculated pairwise distances between cluster centroids (Figure 4A). These distances were used to form a hierarchical clustering, and we calculated the PC-space distance at each branch point (Figure 4B). We found that cluster 3 and 30 cluster together, which together defined the broad mesodermal cell type, even though cluster 3 was primarily composed of juvenile cells and cluster 30 was primarily composed of larval cells (Figure 4B). Larval gut clusters 9 and 35 are closer to larval neural clusters 28 and 26 than to other adult clusters from the juvenile. This is surprising as these cell types arise from separate germ layers, the endoderm and ectoderm. Adult neural clusters 38, 1, 2, and 40 are transcriptionally more similar to adult gut clusters 21, 8, and 27 than they are to larval neural clusters (Figure 4B). These findings supported our predictions based on the observations of the UMAP that some cell types are more similar between life stages than they are between cell types.

**Figure 4.**
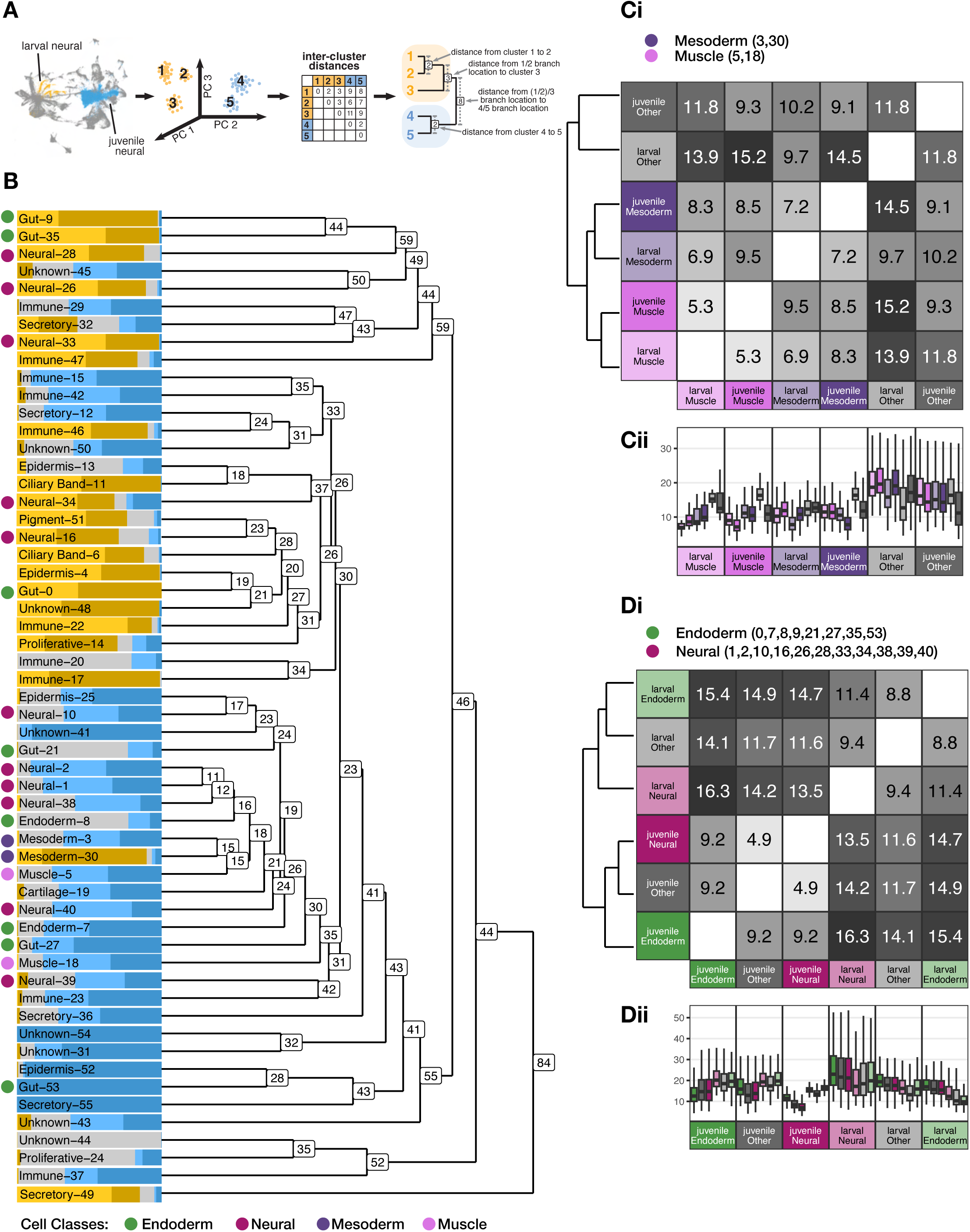
Mesodermal tissues share transcriptional similarity while ectodermal and endoderm tissues reflect life stage divergence. A) Schematic diagram of approach used to accurately quantify similarity or dissimilarity of clusters in 55-dimensional PC space. B) Dendrogram shows a hierarchical clustering of scRNA-seq clusters using the centroid of each cluster in 55-dimensional PC space, with bars colored by cluster life stage composition. Numbers at nodes indicate the distance between the centroids of the two branches merging at that node, as a percent of the largest inter-cluster distance. Colored dots highlight tissues that are further analyzed in Figures Ci, Cii, Di, Dii. C) Comparison of larval mesoderm and larval muscle versus juvenile mesoderm and juvenile muscle, with “Other” denoting all other cells from that life stage. Ci) Heatmap indicates inter-centroid distances as a percent of the largest inter-cluster distance, a larger value means centroids are farther apart while a smaller value means centroids are closer. Cii) Box plots show the distribution from the cells of a group (panel) to the centroids of each other group (bars). D) Comparison of larval neurons and larval endoderm versus juvenile neurons and juvenile endoderm, with “Other” denoting all other cells from that life stage. Di) Heatmap indicates inter-centroid distances as a percent of the largest inter-cluster distance Dii) Box plots show the distribution from the cells of a group (panel) to the centroids of each other group (bars). In box plots, the upper whisker extends from the hinge to the largest value no further than 1.5 * IQR from the hinge (where IQR is the inter-quartile range, or distance between the first and third quartiles). The lower whisker extends from the hinge to the smallest value at most 1.5 * IQR of the hinge.

We next quantified whether transcriptional shifts were larger between cell types in the same life stage or between life stages of the same cell type. To do this, we grouped the cells first by tissues of interest, then split those groups by life stage (all cells outside the tissue of interest are grouped into “Other”). As before, we found the centroids of these groups in transcriptional space and calculated pairwise distances between them, then used these to generate a hierarchical clustering (Figure 4Ci, 4Di). To help evaluate how representative each centroid was of the position of its cells in transcriptional space, we also calculated the distance from individual cells in each group to each group centroid (Figure 4Cii, 4Dii).

Interestingly, we found that larval mesodermally-derived cells, either mesoderm or muscle, are more transcriptionally similar to juvenile mesodermal cells than they are to other larval cell types (Figure 4Ci–4Cii). This is counter to what we observed for many of the endodermally- and ectodermally-derived cell types. For example, larval neurons and larval gut cells are generally more transcriptionally similar than either is to adult neurons or adult gut cells, respectively (Figure 4Di-Dii). This was suggested by the space they occupy in low-dimensional embedding on the UMAP but bolstered by higher-dimensional analysis.

### Genetic makeup of cell type diversity across life history stages

Next, we sought to understand what kind of genes were driving transcriptional differences between larval and juvenile cell types and how those genes were being regulated. We first used GO term analysis to find classes of genes enriched at different life stages in scRNA-seq differential expression data (Figure 5A). These results recapitulated our understanding of hemichordate biology: processes such as cilium organization, assembly, and movement were highest in early and late larva, concomitant with the presence of the circumoral ciliary band and the telotroch, as well as during metamorphosis when the circumoral band has degenerated but the telotroch is still retained. In addition, we observed that GO terms were enriched on a cluster-by-cluster basis (Figure 5B) which also reflected the possible function of these various cell types – for example clusters 6 and 11 were enriched for ciliary components. Ordering the clusters by life stage composition revealed temporal patterns with an enrichment of cellular differentiation and developmental processes starting at metamorphosis. As development proceeds through metamorphosis and into early juvenile and late juvenile stages, we observe an enrichment of GO terms related to morphogenesis like neurogenesis (clusters 16, 1, 2) again reflecting the biological processes occurring at these stages and their underlying cell types (Figure 5B).

**Figure 5.**
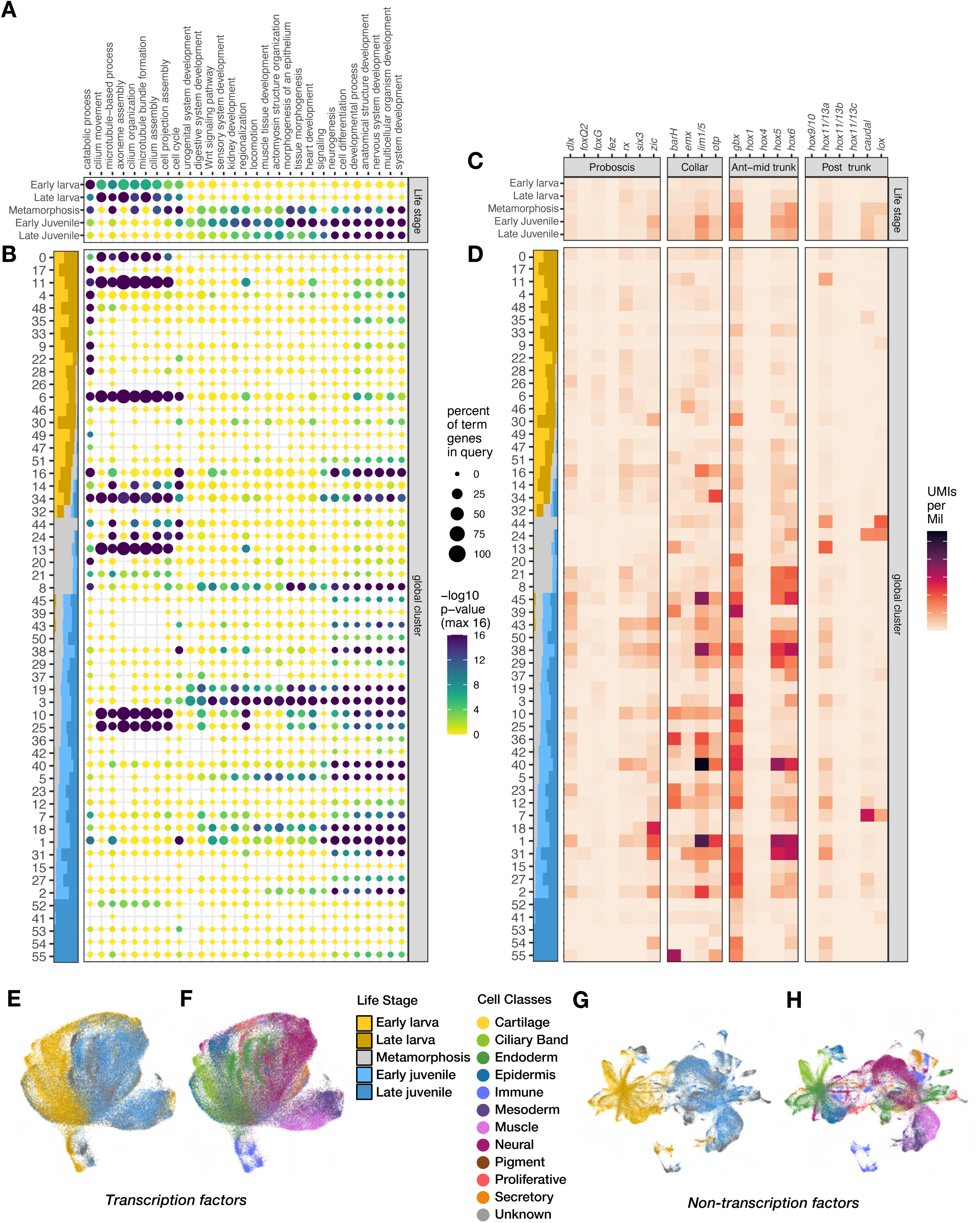
Genetic makeup of cell type diversity across life history stages. A) GO Term analysis of scRNA-seq differential expression data by life stage. B) GO Term analysis of scRNA-seq differential expression data by cluster, ordered by cluster life stage composition per Figure 3D. C) Analysis of anterior-posterior patterning genes in scRNA-seq data by life stage. D) Mean expression of anteroposterior patterning genes in scRNA-seq data by cluster, ordered by cluster composition chart from Figure 3D. E,F) UMAP embedding of scRNA-seq data based on PCA using only transcription factors, colored by life stage (E) or broad cell type (F). G,H) UMAP embedding of scRNA-seq data based on PCA using only non-transcription factors, colored by life stage (G) or broad cell type (H).

Importantly, we also noted enrichment of GO terms associated with the Wnt signaling pathway during metamorphosis and juvenile stages. It has been postulated that in hemichordates the Wnt signaling pathway may be responsible for driving the patterning of posterior Hox-expressing territory (Darras et al., 2018). The anteroposterior patterning program, including the Hox complement (and body patterning more broadly) has previously been described in *S. californicum*, with the larva being mostly “head” or anterior territory, while the trunk region that starts to expand at metamorphosis expresses posterior markers (Gonzalez et al., 2016). We investigated the expression of anteroposterior patterning genes across scRNA-seq, and observed specific enrichment of posterior patterning genes in metamorphosis and juvenile stages compared to larval stages (Figure 5C). More striking, these patterns of gene enrichment also correlated with enrichment of cell types in each life stage (Figure 5D). Indeed, neural clusters, 38, 40, 1, and 2 that are abundant in early and late juvenile stages also highly express genes of the collar and trunk territories, such as *lim1/5*, *gbx*, *hox5*, and *hox6* (Figure 5D).

While these patterning genes may reflect the distribution of these cell types across life stages, we next wanted to examine how different types of genes might explain the contrasts between larval and juvenile cell types. To explore this, we ran a principal component (PC) analysis and UMAP embedding on our scRNA-seq data using either exclusively predicted transcription factors (Figure 5E, 5F) or non-transcription factor genes (Figure 5G, 5H). Both analyses demonstrate some cell separation by life stage (Figure 5E, 5G) and recapitulate separation of the broad cell types (Figure 5F, 5H) we originally identified (Figure 2A). This suggests that both transcription factor and non-transcription factor genes reflect the large effect of physiology and homeostasis between life history stages. This approach demonstrates that regulatory, physiological, and homeostatic programs are all involved in determining cell type composition and transcriptional signatures across disparate life stages. These differences are perhaps linked to the differing ecological demands that body plan morphology must meet as a planktonic larva versus as a mud-dwelling benthic juvenile.

## Discussion

We examined the diversity of cell types in the hemichordate *S. californicum* across five developmental stages: two larval stages, one during metamorphosis, and two juvenile stages. This analysis identified 12 cell classes: cartilage, ciliary band, endoderm, epidermis, immune, mesoderm, muscle, neural, pigment, proliferative, secretory, and unknown. Our findings encompass all major tissue types and include previously unrecognized cell populations.

Some of our findings were unsurprising based on our existing knowledge of hemichordate biology and the loss or gain of cell types associated with loss and gain of structures between life history stages. For example, two ciliary clusters mark the two ciliary bands of the larva, and both the cell clusters and morphological structures are lost following metamorphosis as the benthic adult body plan is formed. Conversely, we observe the appearance of novel clusters associated with the adult body plan, such as the cartilage that is used for structural support of the proboscis and the gill slits.

Our primary objective was to assess how cell type composition changes throughout the organism’s complex life cycle. By characterizing broad cell types, we established new molecular markers to track the distribution of major cell classes across developmental stages.

We considered three explicit hypotheses regarding how cellular composition maps to life history. First, do larval cell types represent a subset of adult cell types, with metamorphosis leading to an expansion of cellular diversity as the organism develops a more complex adult body plan? Second, do larval and adult stages share a substantial portion of their cellular complements, but each stage also possesses unique cell types? Third, are larval and adult body plans composed of distinct cell types, exhibiting minimal overlap in their transcriptional programs?

Surprisingly, our data largely supports the third hypothesis: when life history is overlaid onto the low-dimensional embeddings, there is a marked separation of cell states associated with life history (Figure 3A), and this separation is borne out by higher-dimensional analysis (Figure 4B). Larval cells cluster together to define a larval transcriptional state, largely to the exclusion of juvenile states, with metamorphosis representing an intermediate state. In other words, cells cluster primarily by life history stage rather than by cell type identity. For example, larval neurons and larval gut cells are generally more transcriptionally similar to each other than either is to adult neurons or gut, respectively (Figure 4B, 4Di-ii). This pattern is suggested in UMAP embeddings and confirmed by hierarchical clustering.

Both endoderm and ectoderm-derived cell types follow this stage-specific clustering pattern: larval cells clustering with other larval cells, and adult cells clustering with other adult cells. Mesoderm and muscle are exceptions. These cell types maintain similar transcriptional profiles in both larvae and adults, suggesting that mesoderm undergoes metamorphosis through different mechanisms than the other two germ layers.

### Developmental mechanisms of life history cellular transformation

We propose two hypotheses to explain these observations. First, larval cells may undergo large-scale apoptosis, with adult cells arising from sequestered stem cells. This largely fits with Davidson and colleagues’ “set aside” hypothesis inspired from sea urchin development, where adult structures would presumably originate from sequestered progenitor cells (Peterson et al., 1997) and metamorphosis would represent a total cellular turnover between larvae and adults. The transcriptional state discontinuity observed in endodermally and ectodermally derived cells between larval and adult stages (Figures 3, 4) and the loss of larval ciliary bands at metamorphosis could be consistent with this hypothesis. However, previous work did not reveal extensive apoptosis during metamorphosis, with TUNEL staining restricted to ciliary band cells at the start of metamorphosis (Bump et al., 2022). Furthermore, the extensive carryover of EdU-labeled larval cells through metamorphosis into adulthood that we observed (Figure 3Gi-Hii) is inconsistent with extensive larval cell turnover between life histories.

In the second hypothesis, larval cells may undergo transcriptional reprogramming during metamorphosis via dedifferentiation and redifferentiation, or via transdifferentiation, which has been postulated to play a key role during adult body plan development by transformation of a larva (Arenas-Mena, 2010; Sánchez Alvarado & Yamanaka, 2014). Lineage and functional studies are needed to test for evidence of reprogramming and to differentiate between the possible alternative developmental explanations. However, evidence for transcriptional reprogramming includes cluster 16 (neural cells), which spans all life stages and exhibits spatial continuity in *pburs* expression. Larval *pburs* localizes to the eyespots in the apical organ (Figure 3Ei) and during metamorphosis it is carried over into adult apical ectoderm (Figures 3Eii, Supplemental Figure 5D), suggesting transcriptional programming changes in retained cells.

In contrast, mesoderm does not follow this pattern. Muscle and mesoderm cells retain similar transcriptional states irrespective of life history, as has also been described in a scRNA-seq comparisons between larvae and adult body plans of two different sea urchin species (Paganos et al., 2025). Additionally, similar to the findings of Paganos et al. (2025), some immune cells exhibit overlapping transcriptional states or close interstage relationships (clusters 17, 20, Figure 2A, 3A). In summary, the cellular composition of the transition from larval to adult body plans in the enteropneust *S. californicum* likely involves transcriptional reprogramming of ectodermal and endodermal cells and conservation of the mesodermal cellular program. Future work combining cellular profiling and functional assays will clarify these mechanisms.

### Life history complexities for the construction of cell atlases

Our data suggests that the striking differences in body plan organization of larvae and adults is primarily based on fundamental differences in cell type composition and transcriptional state. As we continue to build a better understanding of metazoan cell type evolution, it is essential that we consider how to integrate life history complexity into a broader synthesis of animal cell type evolution. Currently, the sampling of bilaterian phyla in comparative cell type studies is mostly limited to direct developers or to early larval stages of animals with complex life cycles.

Much of what we understand of the changes in the cellular composition during metamorphosis comes from seminal work in insects. In holometabolous insects many of the larval tissues undergo programmed cell death, but some larval neurons are carried over and reprogrammed during metamorphosis (Gilbert, 2009; Truman et al., 2023). In sea urchins and nemerteans it has been proposed that most larval cells do not persist into adulthood (Davidson et al., 1995; Maslakova, 2010) but this has never been explicitly tested. More broadly in bilaterians, the process of deferred development (Bishop & Hall, 2020), such as temporal shifts in development of the trunk programs (Gonzalez et al., 2016; Martín-Zamora et al., 2023), may help to explain the diversity of animal life cycles. However, the cellular composition differences in relationship to these heterochronies has not been explored in detail. Clearly our understanding of the dynamics of cellular composition in complex life cycles requires further analysis.

Our view of cellular differentiation has been heavily influenced by the findings from model species, and in most cases differentiation in somatic cells is terminal. Some of the earliest branching animals are highly plastic and cell fate changes are very common (Brunet et al., 2021; Sogabe et al., 2019). However, there are also many exceptions to the view that most cells are terminally differentiated – even in vertebrates – and that cellular plasticity is more common than generally acknowledged (Sánchez Alvarado & Yamanaka, 2014).

A growing range of single-cell RNA-seq projects are beginning to provide insights into how cell types change between life history stages (Link et al., 2025; Paganos et al., 2025; Sebé-Pedrós, Chomsky, et al., 2018; Sebé-Pedrós, Saudemont, et al., 2018). Our work demonstrates that specific cellular complements have evolved to support the vastly different ecologies and behaviors of larvae and adults, and that larval cells may have stable rather than terminally differentiated cell states (Sánchez Alvarado & Yamanaka, 2014). This plasticity may be common in species with a similar life history to *S. californicum* but that needs to be explored in further studies. Ultimately, complex life cycles are phylogenetically broadly dispersed and an essential component of animal biodiversity that should be investigated in more depth and integrated into our understanding of cell type differentiation and evolution.

## Methods

### Collecting, spawning, and larval rearing

Adult *Schizocardium californicum* were collected in Morro Bay State Park, California, in a mudflat located at 35°20′56.7”N 120°50′35.6”W with appropriate state permitting. Animals were spawned as described in Gonzalez et al. 2018 with individual females transferred in bowls of filtered seawater (FSW) and placed in an illuminated incubator at 24–26°C. Once embryos hatched, larvae were transferred to 1 gallon glass jars with continuous stirring and fed with a 1:1 mix of *Dunaliella tertiolecta* and *Rhodomonas lens*. Every two to four days, containers were washed, water was replaced with clean FSW, and fresh algae was added. To grow larger numbers of animals through metamorphosis, some larvae were placed on a continuous flow-through system by being transferred into diffusion tubes (Patry et al., 2020). Once animals began metamorphosis, they were transferred into glass bowls with terrarium sand.

### Single-cell sequencing

#### Cell preparation

Early and late juveniles were first diced into fine pieces with a razor blade on a petri dish. All stages were suspended in 1 mg/mL Dispase II (Sigma D4693) in 3.3× PBS, 2% Fetal Calf Serum, and 20 mM HEPES. Cells were pipetted intermittently with Pasteur pipettes passed over a flame to create narrower openings. Cells were then passed through a 100 μm mesh and centrifuged at 4°C, 150 g, for 5 minutes with slow braking. Cells were resuspended in fresh media and passed through a 40 μm strainer on ice. Cell viability was checked with Hoechst and counted on a hemocytometer. Importantly, cells were kept in a 3.3× PBS solution to maintain osmolarity to seawater.

#### Sequencing

Cells were loaded into the 10x Chromium Controller for cell capture and libraries were processed after SPRI size-selection with the 10x Genomics V2.0 workflow. Final libraries were analyzed on an Agilent Bioanalyzer and sequenced using an Illumina Novaseq.

#### Data processing

Once reads were acquired, they were demultiplexed, mapped, and subsequently converted into count matrices using 10x Genomics Cell Ranger 3.0.0 (G. X. Y. Zheng et al., 2017). Cell quality filtering, clustering, and dimensionality reduction was done using Seurat v3 in R (Hao et al., 2021; Satija et al., 2015; Stuart et al., 2019). Cells were filtered to those with at least 500 and less than 57,000 unique molecular identifiers (UMIs) and >90 and no more than 980 genes. We normalized each batch’s gene expression using Seurat’s SCTransform, then combined the batches using reciprocal principal component analysis (rPCA) integration (Stuart et al., 2019). This technique was chosen both for its efficiency on large datasets, and because rPCA integration by batch resulted in good overlap of the same life stage from different batches without forcing overlap of different life stages. After this we performed principal component analysis (PCA) on the combined dataset with 55 principal components (PCs), generated a shared nearest neighbor graph, used clustering to identify 56 cell clusters, and determined which genes are most differentially expressed across the dataset.

### Alternative integration approaches

To integrate data sets and test for batch effects we used the Portal algorithm (Zhao et al., 2022). Each batch was read in from the Seurat file and integrated incrementally. The datasets were preprocessed and integrated with default parameters and 1000 training steps. We recovered the harmonized expression matrix in log-normalized level. We then processed this expression matrix with the Self-Assembling-Manifold (SAM) algorithm (Tarashansky et al., 2019), with default parameters except for without log normalization or filtering genes. Under these parameters we recovered no significant batch effects.

To integrate data sets and test for batch effects we ran “Harmony” on our principle component embeddings which projects cells into a shared embedding where cells group by cell type rather than dataset-specific conditions (Korsunsky et al., 2019). Independently, we took the cell count matrices and used the Portal algorithm where each batch was read in and integrated incrementally (Zhao et al., 2022). The datasets were preprocessed and integrated with default parameters and 1000 training steps. We recovered the harmonized expression matrix in log-normalized level. We then processed this expression matrix with the Self-Assembling-Manifold (SAM) algorithm (Tarashansky et al., 2019) with default parameters except for without log normalization or filtering genes. Under both approaches we recovered distinct differences between life history stage and no significant batch effects.

### GO Analysis

We first performed differential gene expression of positive markers as calculated across the relevant groups (cluster, life stage) using Seurat’s FindMarkers on the SCTransform-normalized single-cell expression data. We then filtered these results to genes with 1) average log2 fold-change of at least 1.25; 2) adjusted p-value less than 0.0001; 3) which had a reciprocal best BLASTp hit between *Schizocardium californicum* and *Mus musculus* GRCm39 predicted peptide sequences. These differential expression results were then sorted by log2 fold-change and fed into gprofiler2’s gost function with organism=“mmusculus” and ordered_query=TRUE(Kolberg et al., 2023). GO Terms were manually curated for plotting.

### High-dimensional analysis

To perform high-dimensional analysis of overall transcriptional similarity of clusters and cell types, the centroid of each cluster was calculated in 55-dimensional PC space, inter-cluster distances were found for all pairs of clusters, and these distances were scaled as the percentage of the largest inter-cluster distance. These distances were then used to build a hierarchical clustering of all clusters in the dataset. For more detailed analysis, cells were grouped by the combination of cell type and life stage, group centroids and inter-group distances calculated, and these distances scaled by the same factor as the inter-cluster distances to facilitate direct comparison. Then the distribution of distances from each group’s individual cells to each group centroid was determined to assess the accuracy of inter-centroid distances.

### Ortholog identification

Gene orthology and naming was determined by collecting sequences of interest from related species and then building gene trees. Sequences were aligned with MUSCLE (Edgar, 2004) and trees were calculated with Bayesian inference trees using MrBayes version 3.1.2 (Huelsenbeck & Ronquist, 2001) in 1,000,000 generations with sampling of trees every 100 generations and a burn-in period of 25%.

### In situ HCR

#### Probe design and preparation

Complementary DNA sequences specific to genes of interest were submitted to the in situ probe generator from the Ozpolat Lab (Kuehn et al., 2021). The sequences generated by the software were used to order DNA oligo pools (50 μmol DNA oPools™ Oligo Pools) from Integrated DNA Technologies, resuspended to 1 μmol/μL in 50 mM Tris buffer, pH 7.5. HCR amplifiers with fluorophores B1-Alexa Fluor-546, B2-Alexa Fluor-488, and B3-Alexa Fluor-647 were ordered from Molecular Instruments, Inc.

#### HCR protocol

HCRs were performed based on Choi et al., 2018 and the Hybridization Chain Reaction (HCR) in situ Protocol from the Patel Lab (Bruce et al., 2021; Choi et al., 2018). Animals were fixed with 3.7% paraformaldehyde in MOPS fix buffer (0.1 M MOPS, 0.5 M NaCl, 2 mM EGTA, 1 mM MgCl2, 1× PBS) overnight at 4°C. Fixed samples were then dehydrated in methanol and stored at −20°C. Samples were rehydrated in a gradual series of PBS containing 0.1% Tween-20 (PBST). They were permeabilized in detergent solution (1.0% SDS, 0.5% Tween-20, 150 mM NaCl, 1 mM EDTA (pH 8), 50 mM Tris-HCl at pH 7.5) for 30 minutes. The samples were then extensively washed in PBST, before being pre-hybridized in hybridization buffer (Molecular Instruments) for 1 h at 37°C. The probes were then added to the hybridization buffer at a final concentration of 0.05 μM and the samples were allowed to hybridize at 37°C overnight under gentle agitation. Following hybridization, the samples were washed 4 times for 15 minutes in probe wash buffer (Molecular Instruments) at 37°C and then in 5× SSCT at room temperature. They were then pre-amplified in amplification buffer (Molecular Instruments) for 30 minutes. Amplifiers were separately incubated at 95°C for 90 seconds and then pooled together before being added to the amplification buffer at a final concentration of 60 nM. Amplification was performed overnight, and the samples were subsequently washed 4 times for 30 minutes in 5× SSCT and incubated in 5× SSCT containing 1:1000 Hoechst overnight.

### Pulse-chase EdU labeling

Labeling and detection of proliferating cells were performed using the Click-iT Plus EdU 488 Imaging Kit (Life Technologies), using the standard protocol with the following modifications. Larvae were cultured in FSW supplemented with 10 μM EdU diluted from a 10 mM stock in DMSO. Animals were pulsed for 48 hours with EdU then either fixed with 3.7% paraformaldehyde in MOPS fix buffer (0.1 M MOPS, 0.5 M NaCl, 2 mM EGTA, 1 mM MgCl2, 1× PBS) overnight at 4°C or transferred to fresh FSW and allowed to grow and were fixed at either 43 or 26 days later when they had completed metamorphosis. For detection of EdU incorporation, labeled tissue was transferred to a solution of PBS and the detection was performed following the manufacturer’s protocol with an increased permeabilization time in 0.5% Triton X-100 in PBS of 40 minutes and an increased detection time of 45 minutes.

### Imaging

Samples were mounted in PBS (larva) or fructose (juveniles) between two glass coverslips raised slightly with small clay feet. Imaging was performed on a Zeiss LSM 700. For large samples, stacks were stitched together using the Tile tool (Zen2 software, Zeiss).

## Acknowledgements

We would like to thank the staff of the Hopkins Marine Station and the staff of Morro Bay State Park, in particular Vince Cicero, John Sayers, and Katie Drexhage, for facilitating our collections. We would like to thank Norma Neff from Biohub for supporting the development of genomic resources for *Schizocardium*. We thank members of the Lowe Lab, specifically Auston Rutledge who reared many *Schizocardium* larvae and was an invaluable partner in collections. We also thank Greg Wray, Bo Wang, Dominique Bergmann, Stephen Palumbi, Lowe Lab members Nat Clarke, Paul Minor, Veronica Pagowski, José Andrade-Lopez, and Srivastava Lab members Mansi Srivastava, Katy Loubet-Senear, Amber Rock, Carlos Rivera-Lopez and Catie Breen for their helpful discussions. We finally thank previous Lowe Lab member Paul Gonzalez for his pioneering work on *Schizocardium*. This work was supported by a collaborative CZ Biohub Intercampus Research Award to C.J.L and a NSF predoctoral fellowship (DGE– 1147470), the Myers Trust Award, and Haderlie Memorial Award to P.B.

## Data availability

The sequencing raw reads and data generated in this study have been deposited in the NCBI BioProject database under the accession code PRJNA1297089.

## Supplemental Figure Legends

**Supplemental Figure 1.**
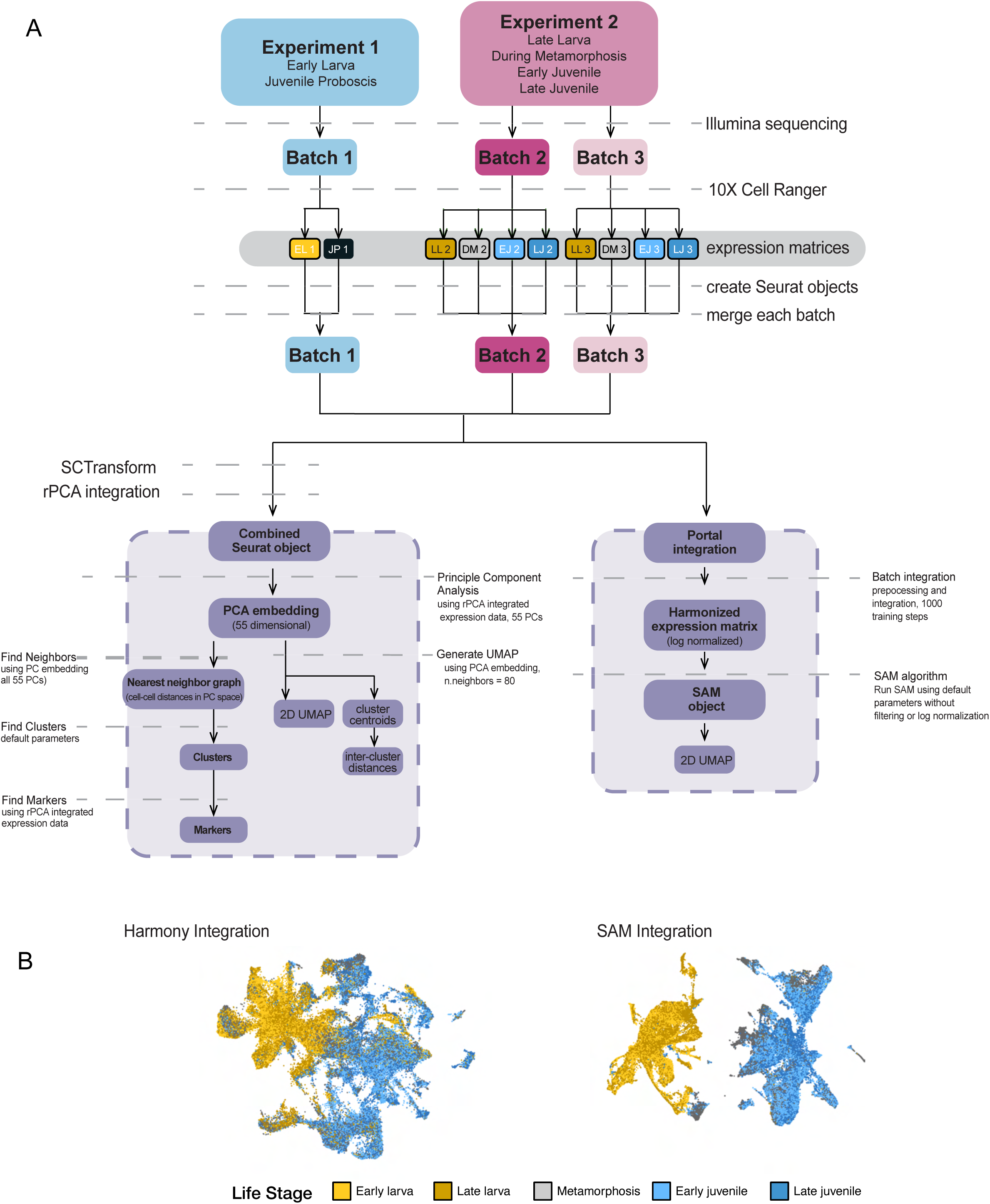
A) Flowchart of single-cell analysis including alternative integration approaches. B) UMAP embedding of alternative integration approaches, “Harmony” and “Portal” plotted by life stage.

**Supplemental Figure 2.**
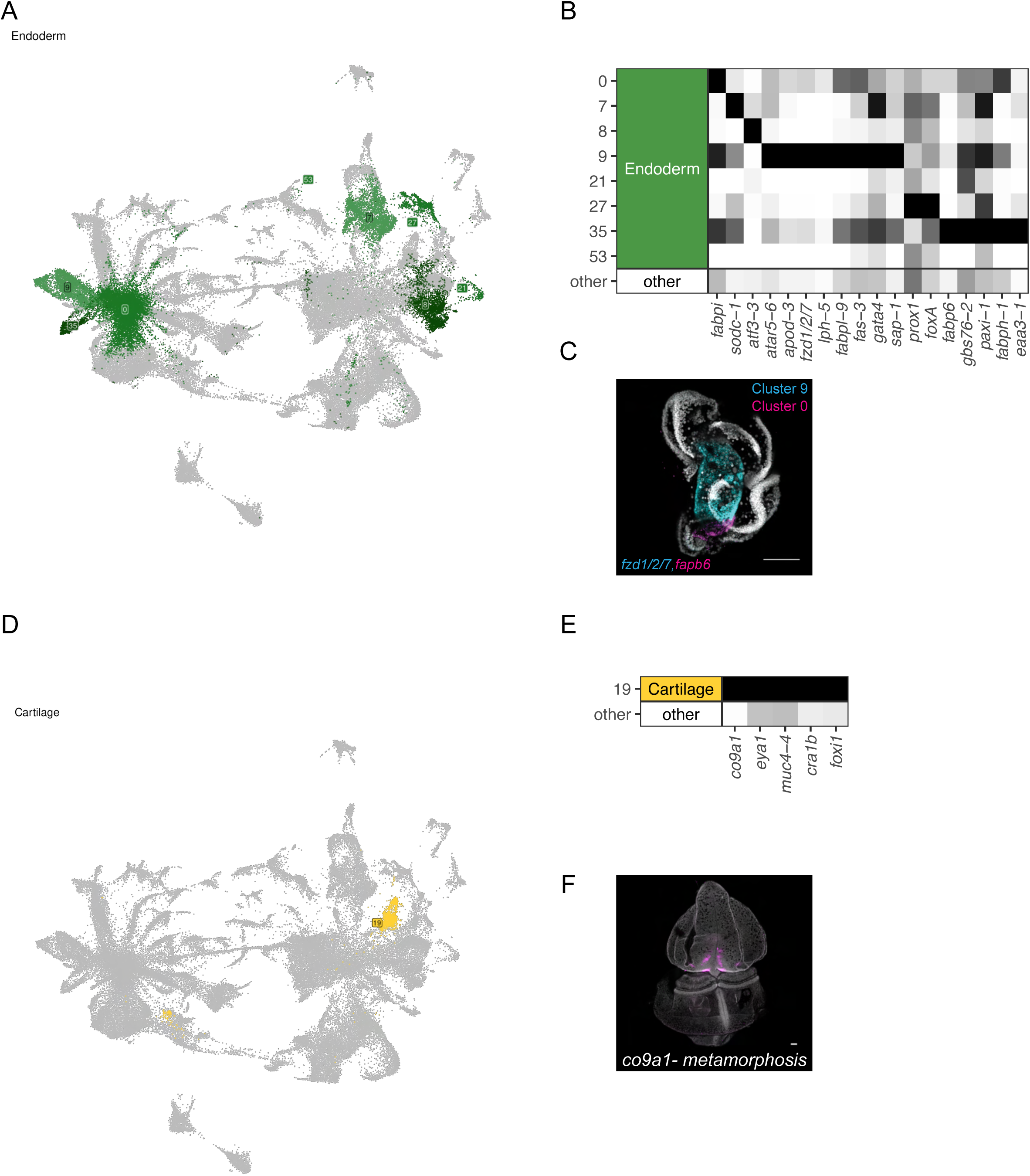
A) Endoderm clusters (*0, 7, 8, 9, 21, 27, 35, 53)* plotted in global UMAP embedding. B) Endoderm cluster expression of marker genes. C) HCR expression in early larva of *fzd1/2/7* in cyan, *fabp6* in magenta, nuclei in grey, scale bars are 100 µm. D) Cartilage cluster (*19)* plotted in global UMAP embedding. E) Cartilage cluster expression of marker genes. F) HCR expression during metamorphosis of *co9a1* in magenta, nuclei grey, scale bar 100 µm.

**Supplemental Figure 3.**
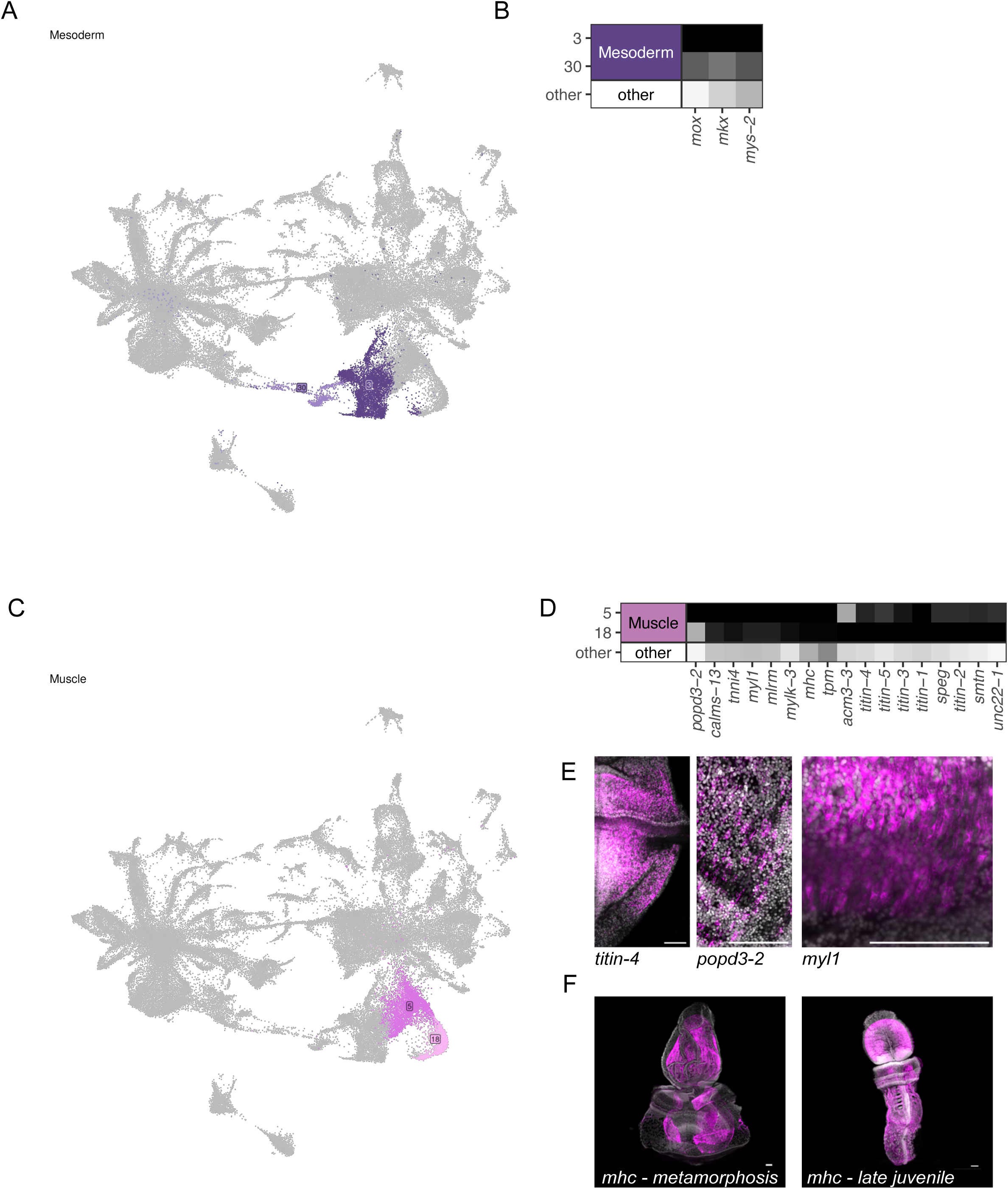
A) Mesoderm clusters (*3, 30)* plotted in global UMAP embedding. B) Mesoderm cluster expression of marker genes. C) Muscle clusters (*5, 18)* plotted in global UMAP embedding. D) Muscle cluster expression of marker genes. E) HCR expression of muscle markers *titin-4, popd3-2, myl1* in magenta, nuclei in grey, scale bars are 100 µm. F) HCR expression during metamorphosis and in late juvenile of *mhc* in magenta, nuclei in grey, scale bars are 100 µm.

**Supplemental Figure 4.**
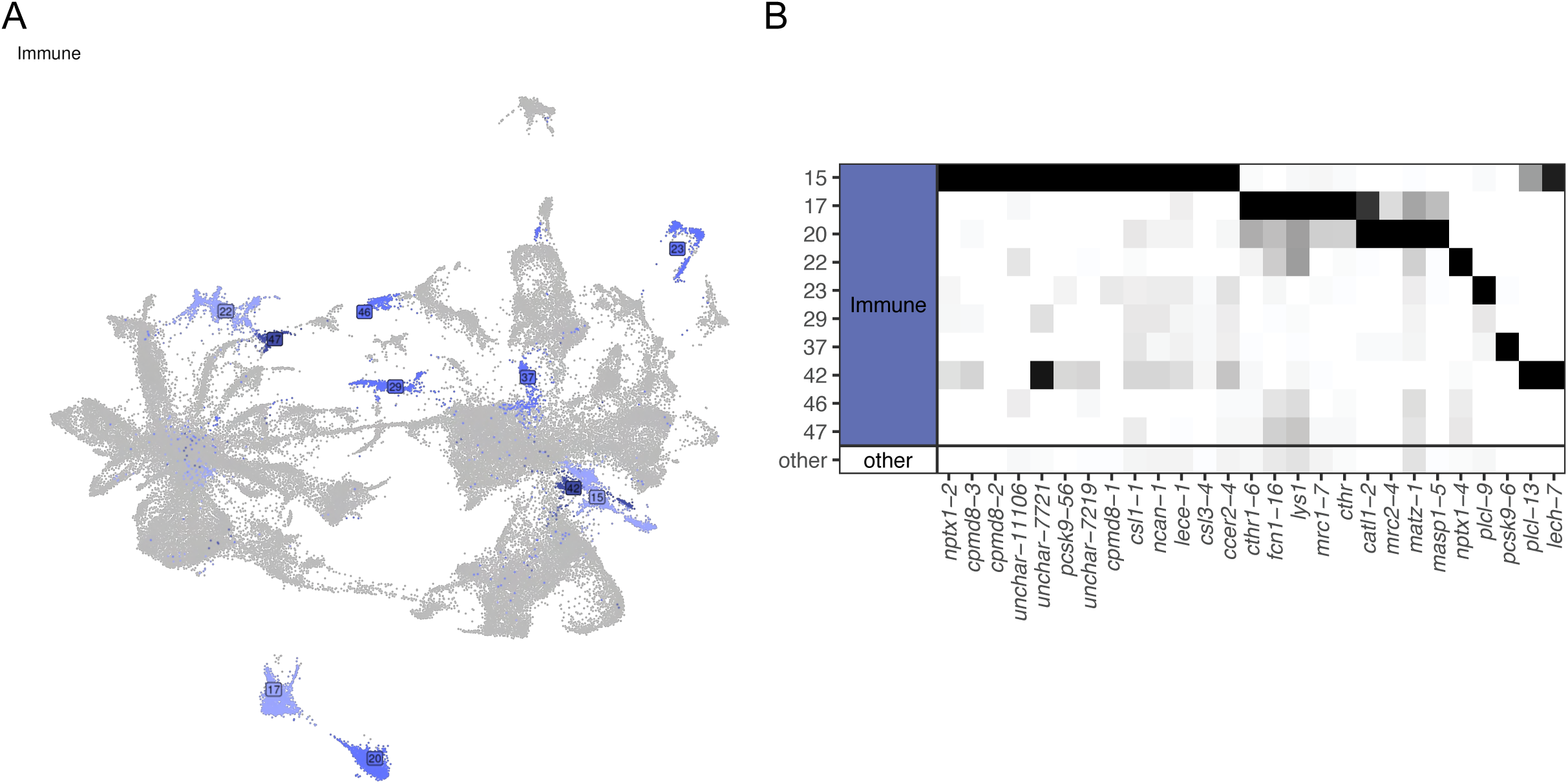
A) Immune clusters (*15, 17, 20, 22, 23, 29, 37, 42, 46, 47)* plotted in global UMAP embedding. B) Immune cluster expression of marker genes.

**Supplemental Figure 5.**
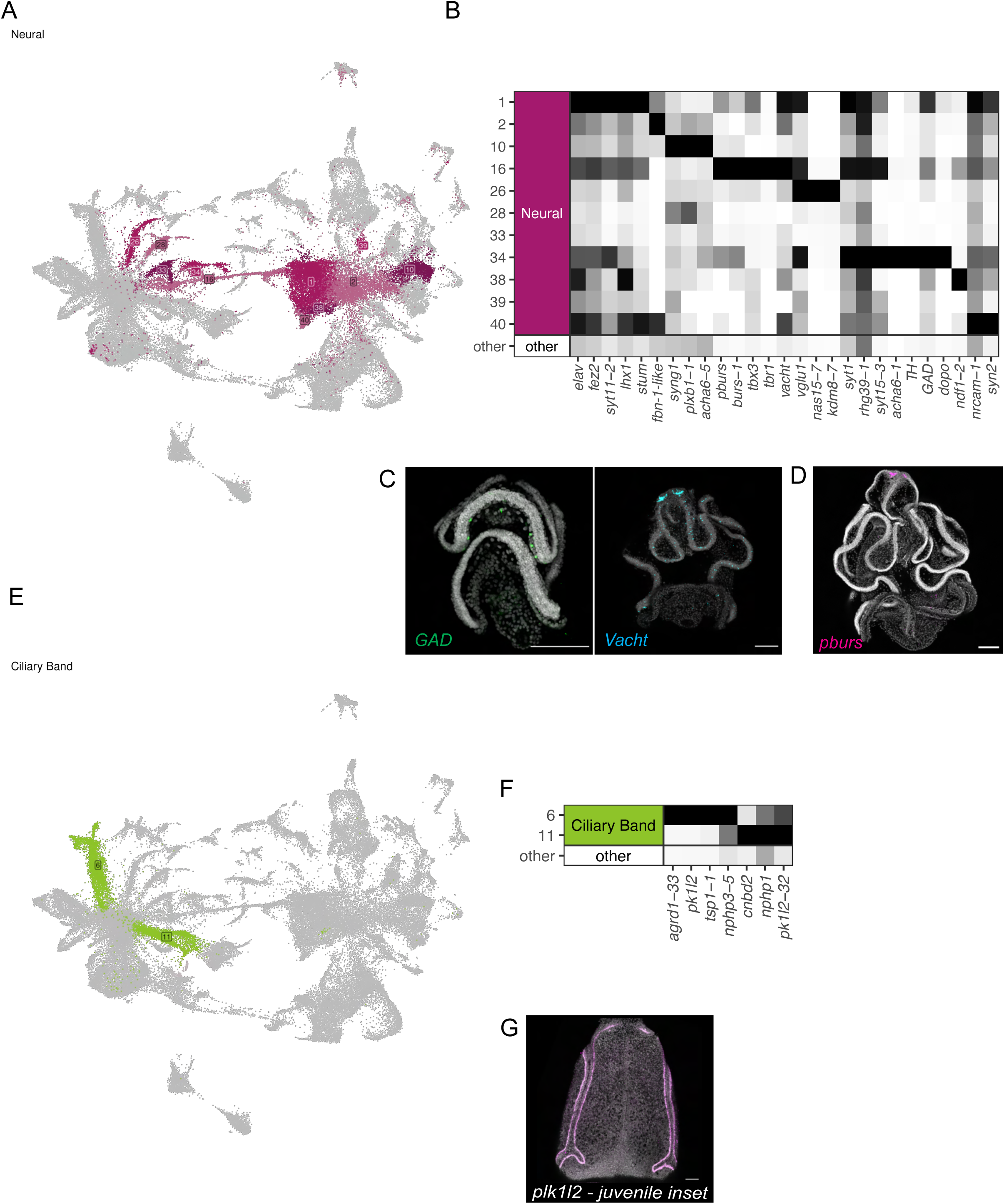
A) Neural clusters (*1, 2, 10, 16, 26, 28, 33, 34, 38, 39, 40)* plotted in global UMAP embedding. B) Neural cluster expression of marker genes. C) HCR expression of neural markers *GAD* (green), *VAChT* (cyan), nuclei in grey, scale bars are 100 µm. D) HCR expression in late larva of *pburs* (magena), nuclei in grey, scale bars are 100 µm. E) Ciliary clusters (*6, 11)* plotted in global UMAP embedding. F) Ciliary cluster expression of marker genes. G) HCR expression in early juvenile proboscis inset of *pk1l2* in magenta, nuclei in grey, scale bars are 100 µm.

**Supplemental Figure 6.**
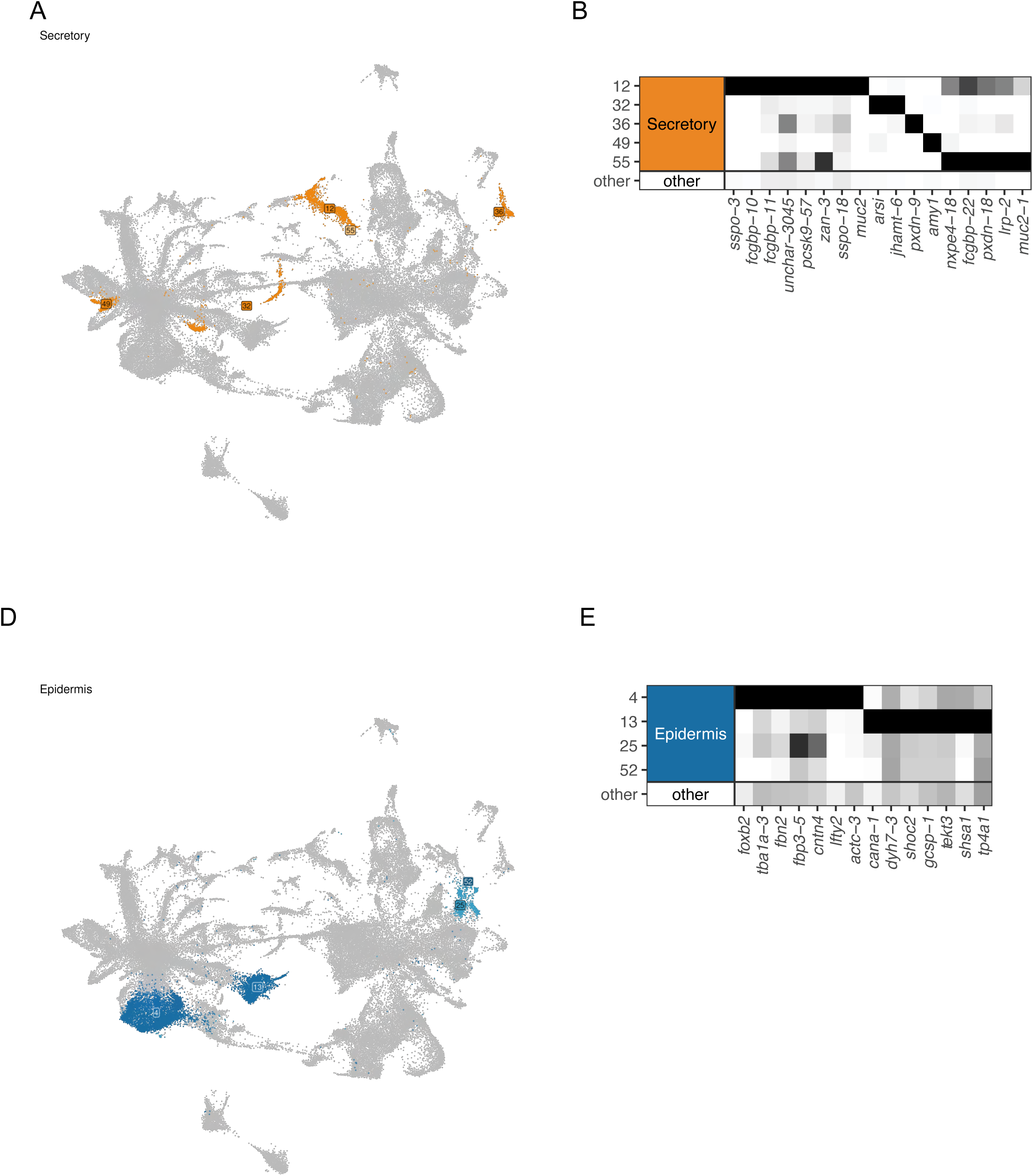
A) Secretory clusters (*1, 2, 10, 16, 26, 28, 33, 34, 38, 39, 40)* plotted in global UMAP embedding. B) Secretory cluster expression of marker genes. C) Epidermis clusters (*4, 13, 25, 52)* plotted in global UMAP embedding. D) Epidermis cluster expression of marker genes.

**Supplemental Figure 7.**
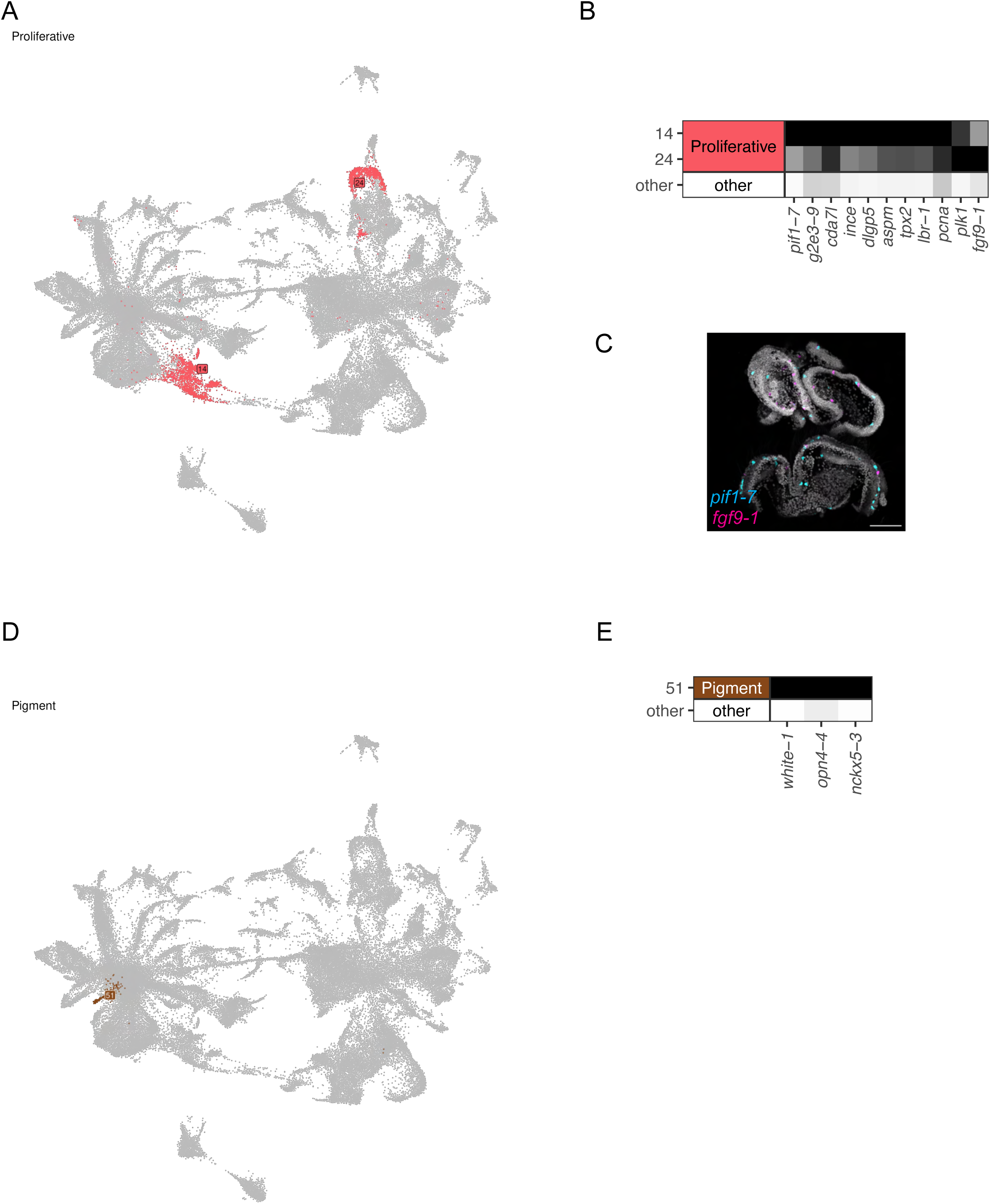
A) Proliferative clusters (*14, 24)* plotted in global UMAP embedding. B) Proliferative cluster expression of marker genes. C) HCR expression in early larva of *pif1-7* in cyan, *fgf9-1* in magenta, nuclei in grey, scale bars are 100 µm. D) Pigment cluster (*51)* plotted in global UMAP embedding. E) Pigment cluster expression of marker genes.

**Supplemental Figure 8.**
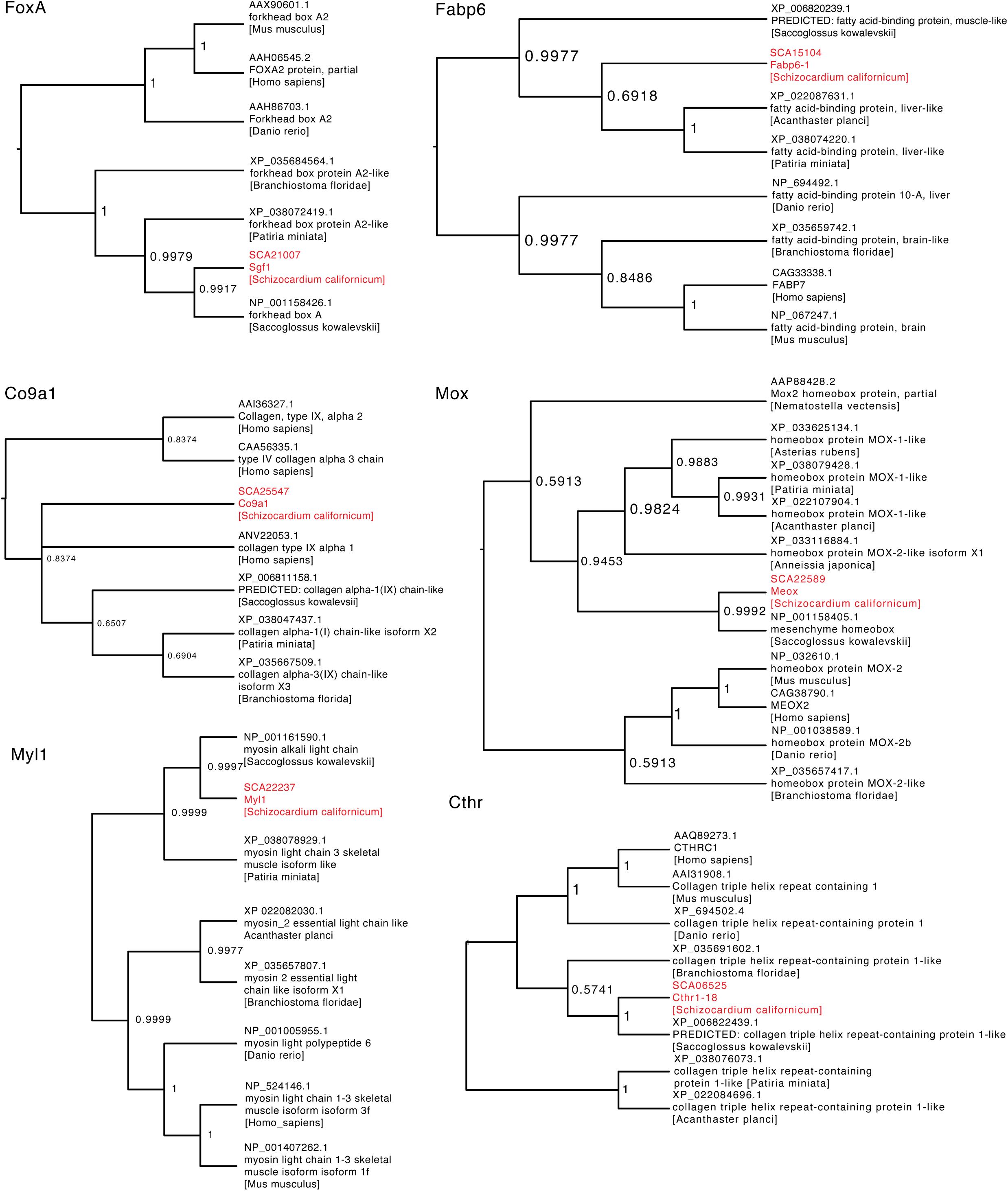
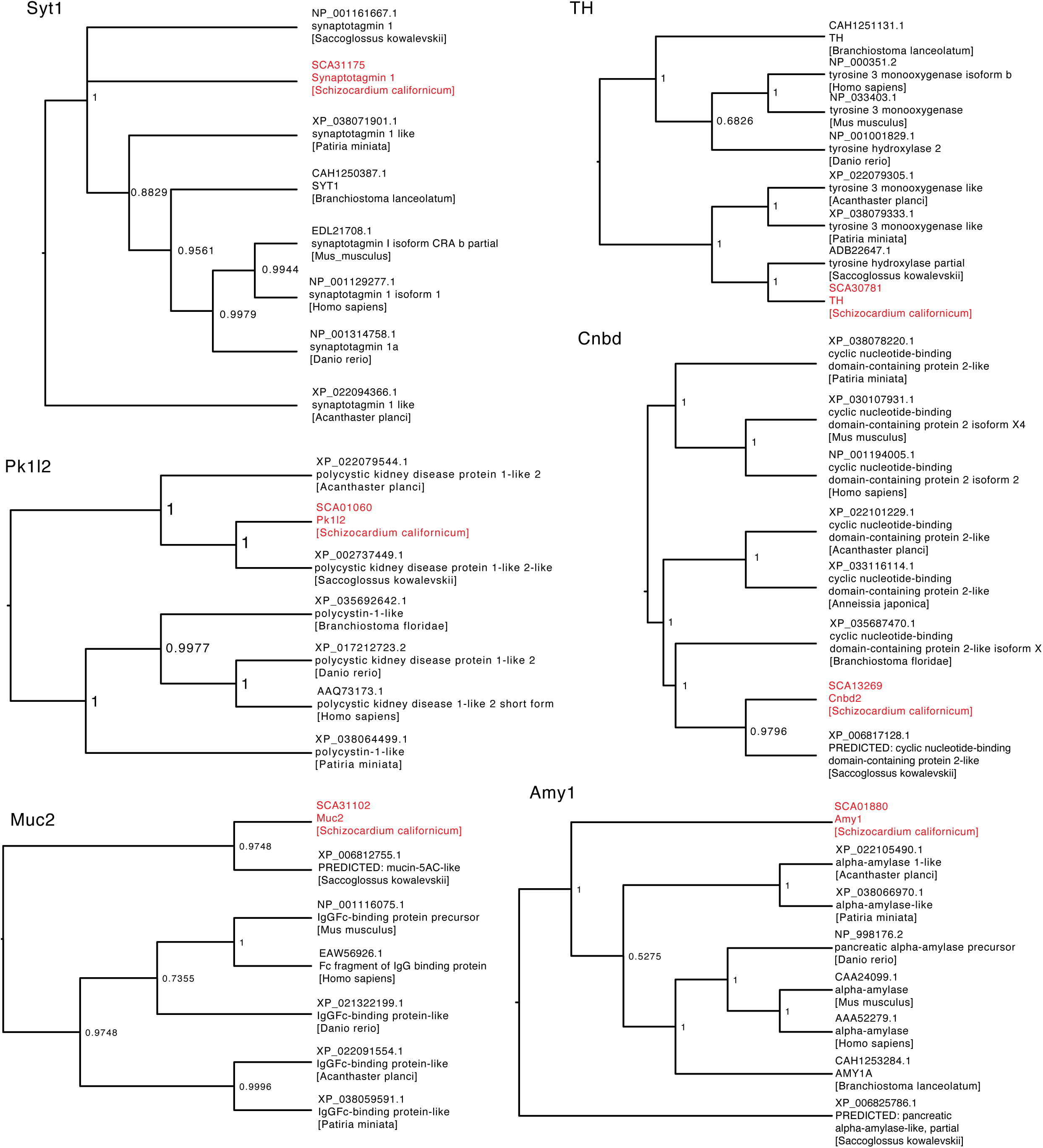
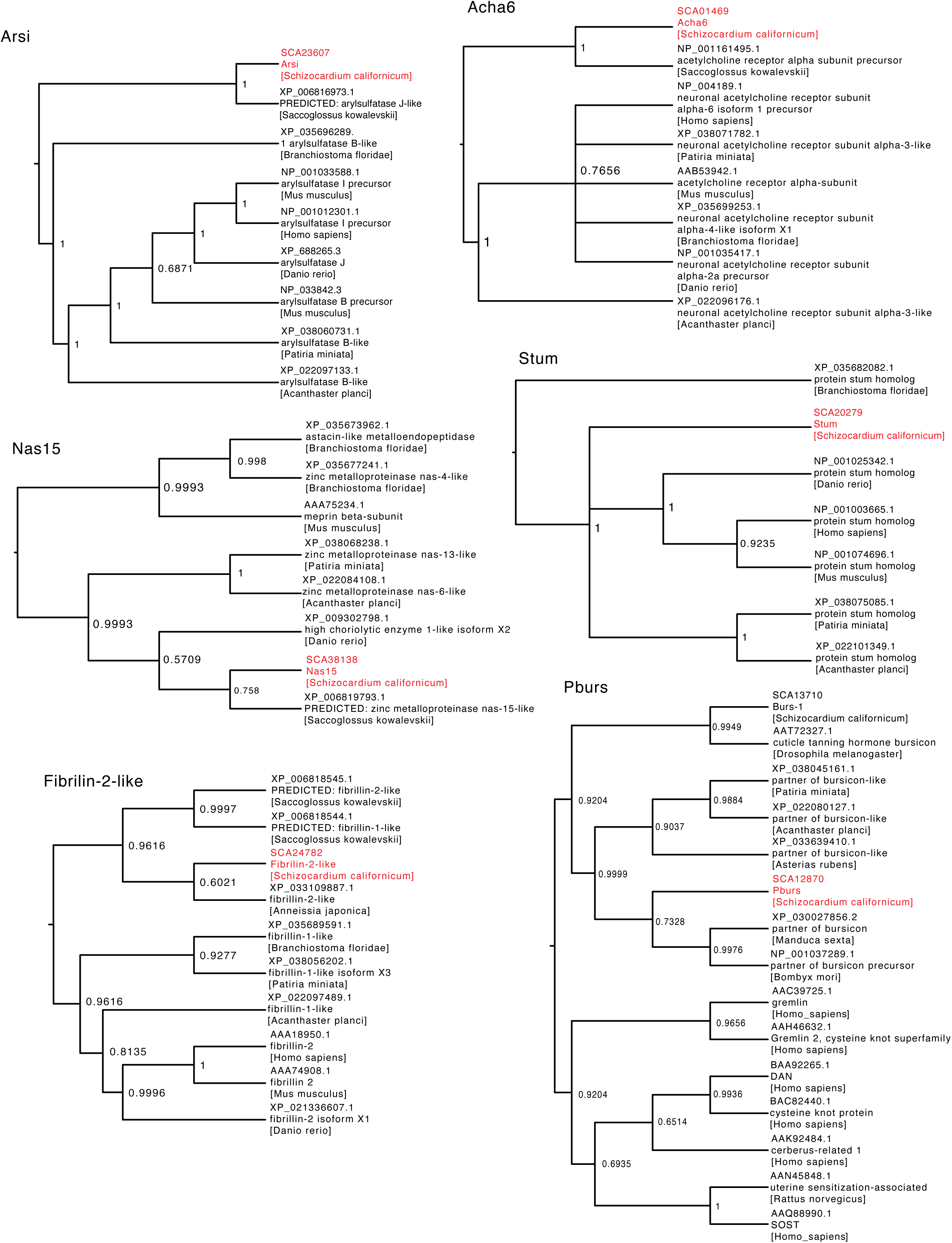
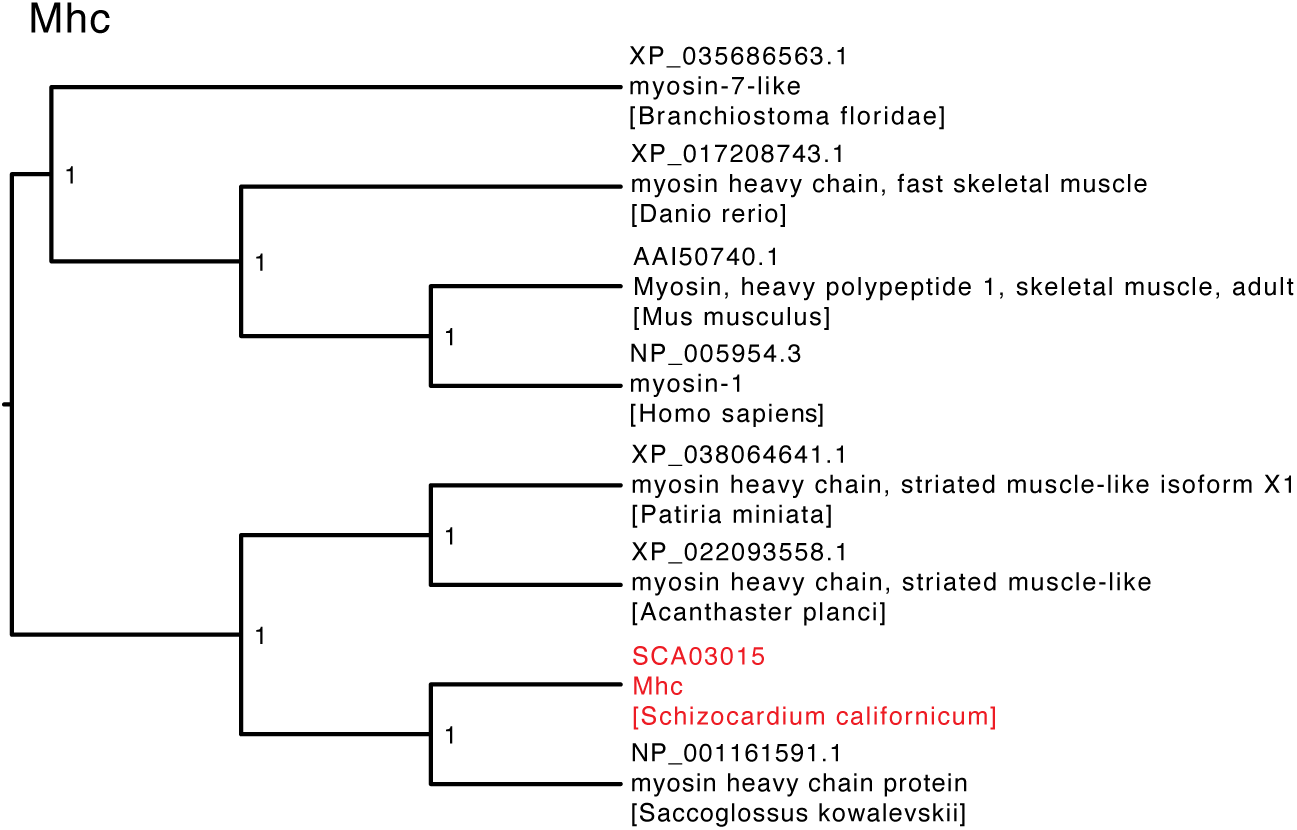
Phylogenetic trees of *Schizocardium californicum* orthologs.

## Supplemental Tables

**Supplemental Table 1.** scRNA-seq statistics for 56 clusters identified and cell class annotation

**Supplemental Table 2.** Cell cluster markers obtained by analysis of differential expression (log fold change and p-value) and specificity (how frequently a marker appears).

**Supplemental Table 3.** Cell class markers including gene nomenclature

**Supplemental Table 4.** HCR probe sequences with corresponding amplifier design (B1, B2, or B3).

## References

Agassiz, A. (1873). The History of Balanoglossus and Tornaria. Memoirs of the American Academy of Arts and Sciences, 9(2), 421–436. JSTOR. 10.2307/25058009

Andrade López, J. M., Pani, A. M., Wu, M., Gerhart, J., & Lowe, C. J. (2023). Molecular characterization of nervous system organization in the hemichordate acorn worm Saccoglossus kowalevskii. PLOS Biology, 21(9), e3002242. 10.1371/journal.pbio.3002242

Arenas-Mena, C. (2010). Indirect development, transdifferentiation and the macroregulatory evolution of metazoans. Philosophical Transactions of the Royal Society B: Biological Sciences, 365(1540), 653–669. 10.1098/rstb.2009.0253

Axelrod, J. (1972). Dopamine-β-hydroxylase: Regulation of Its Synthesis and Release from Nerve Terminals. Pharmacological Reviews, 24(2), 233–243.

Balser, E. J., & Ruppert, E. E. (1990). Structure, Ultrastructure, and Function of the Preoral Heart-Kidney in *Saccoglossus kowalevskii* (Hemichordata, Enteropneusta) Including New Data on the Stomochord. Acta Zoologica, 71(4), 235–249. 10.1111/j.1463-6395.1990.tb01082.x

Berger, C., Renner, S., Lüer, K., & Technau, G. M. (2007). The commonly used marker ELAV is transiently expressed in neuroblasts and glial cells in the *Drosophila* embryonic CNS. Developmental Dynamics, 236(12), 3562–3568. 10.1002/dvdy.21372

Betz, H. (1990). Ligand-gated ion channels in the brain: The amino acid receptor superfamily. Neuron, 5(4), 383–392. 10.1016/0896-6273(90)90077-s

Bishop, C., & Hall, B. (2020). Deferring Development: Setting Aside Cells for Future Use in Development and Evolution. CRC Press/Taylor & Francis Group.

Bruce, H., Jerz, G., Kelly, S., McCarthy, J., Pomerantz, A., Senevirathne, G., Sherrard, A., Sun, D., Wolff, C., & Patel, N. (2021). Hybridization Chain Reaction (HCR) In Situ Protocol. Protocols.Io. 10.17504/protocols.io.bunznvf6

Brunet, T., Albert, M., Roman, W., Coyle, M. C., Spitzer, D. C., & King, N. (2021). A flagellate-to-amoeboid switch in the closest living relatives of animals. eLife, 10, e61037. 10.7554/eLife.61037

Bullock, T. H. (1945). The Anatomical Organization of the Nervous System of Enteropneusta. Journal of Cell Science, s2-86(341), 55–111. 10.1242/jcs.s2-86.341.55

Bump, P., Khariton, M., Stubbert, C., Moyen, N. E., Yan, J., Wang, B., & Lowe, C. J. (2022). Comparisons of cell proliferation and cell death from tornaria larva to juvenile worm in the hemichordate *Schizocardium californicum*. EvoDevo, 13(1), 13. 10.1186/s13227-022-00198-1

Cameron, C. B. (2002). Particle Retention and Flow in the Pharynx of the Enteropneust Worm *Harrimania planktophilus*: The Filter-Feeding Pharynx May Have Evolved Before the Chordates. The Biological Bulletin, 202(2), 192–200. 10.2307/1543655

Cameron, C. B., & Perez, M. (2012). Spengelidae (Hemichordate: Enteropnuesta) from the Eastern Pacific including a new species, *Schizocardium californicum*, from California. Zootaxa, 3569, 79–88.

Chernousov, M. A., Baylor, K., Stahl, R. C., Stecker, M. M., Sakai, L. Y., Lee-Arteaga, S., Ramirez, F., & Carey, D. J. (2010). Fibrillin-2 is dispensable for peripheral nerve development, myelination and regeneration. Matrix Biology : Journal of the International Society for Matrix Biology, 29(5), 357–368. 10.1016/j.matbio.2010.02.006

Choi, H. M. T., Schwarzkopf, M., Fornace, M. E., Acharya, A., Artavanis, G., Stegmaier, J., Cunha, A., & Pierce, N. A. (2018). Third-generation *in situ* hybridization chain reaction: Multiplexed, quantitative, sensitive, versatile, robust. Development, 145(12), dev165753. 10.1242/dev.165753

Cunningham, D., & Casey, E. S. (2014). Spatiotemporal development of the embryonic nervous system of *Saccoglossus kowalevskii*. Developmental Biology, 386(1), 252–263. 10.1016/j.ydbio.2013.12.001

Darras, S., Fritzenwanker, J. H., Uhlinger, K. R., Farrelly, E., Pani, A. M., Hurley, I. A., Norris, R. P., Osovitz, M., Terasaki, M., Wu, M., Aronowicz, J., Kirschner, M., Gerhart, J. C., & Lowe, C. J. (2018). Anteroposterior axis patterning by early canonical Wnt signaling during hemichordate development. PLoS Biology, 16(1). 10.1371/journal.pbio.2003698

Darszon, A., Guerrero, A., Galindo, B. E., Nishigaki, T., & Wood, C. D. (2008). Sperm-activating peptides in the regulation of ion fluxes, signal transduction and motility. The International Journal of Developmental Biology, 52(5–6), 595–606. 10.1387/ijdb.072550ad

Das, B., Cai, L., Carter, M. G., Piao, Y.-L., Sharov, A. A., Ko, M. S. H., & Brown, D. D. (2006). Gene expression changes at metamorphosis induced by thyroid hormone in *Xenopus laevis* tadpoles. Developmental Biology, 291(2), 342–355. 10.1016/j.ydbio.2005.12.032

Davidson, E. H., Peterson, K. J., & Cameron, R. A. (1995). Origin of Bilaterian Body Plans: Evolution of Developmental Regulatory Mechanisms. Science, 270(5240), 1319–1325. 10.1126/science.270.5240.1319

Desai, B. S., Chadha, A., & Cook, B. (2014). The stum Gene Is Essential for Mechanical Sensing in Proprioceptive Neurons. Science, 343(6176), 1256–1259. 10.1126/science.1247761

Edgar, R. C. (2004). MUSCLE: multiple sequence alignment with high accuracy and high throughput. Nucleic Acids Research, 32(5), 1792–1797. PubMed. 10.1093/nar/gkh340

Formery, L., & Lowe, C. J. (2023). Integrating Complex Life Cycles in Comparative Developmental Biology.

Foster, S., Oulhen, N., Fresques, T., Zaki, H., & Wessel, G. (2022). Single-cell RNA-sequencing analysis of early sea star development. Development, 149(22), dev200982. 10.1242/dev.200982

Fritzenwanker, J. H., Gerhart, J., Freeman, R. M., & Lowe, C. J. (2014). The Fox/Forkhead transcription factor family of the hemichordate *Saccoglossus kowalevskii*. EvoDevo, 5(1), 17. 10.1186/2041-9139-5-17

Garstang, W. (1939). Spolia Bermudiana. II. The ciliary feeding mechanism of Tornaria. Q. Jl Microsc. Sci., 81, 347–366.

Gilbert, L. I. (2009). Insect development: Morphogenesis, molting and metamorphosis. Academic Press.

Gonzalez, P., & Cameron, C. (2009). The gill slits and pre-oral ciliary organ of *Protoglossus* (Hemichordata: Enteropneusta) are filter-feeding structures. Biological Journal of the Linnean Society, 98(4), 898–906. 10.1111/j.1095-8312.2009.01332.x

Gonzalez, P., Jiang, J. Z., & Lowe, C. J. (2018). The development and metamorphosis of the indirect developing acorn worm *Schizocardium californicuma* (Enteropneusta: Spengelidae). Frontiers in Zoology, 1–24.

Gonzalez, P., Uhlinger, K. R., & Lowe, C. J. (2016). The Adult Body Plan of Indirect Developing Hemichordates Develops by Adding a Hox-Patterned Trunk to an Anterior Larval Territory. Current Biology, 27(1), 1–9. 10.1016/j.cub.2016.10.047

Green, S. A., Norris, R. P., Terasaki, M., & Lowe, C. J. (2013). FGF signaling induces mesoderm in the hemichordate *Saccoglossus kowalevskii*. Development, 140(5), 1024– 1033. 10.1242/dev.083790

Hadfield, M. G., Carpizo-Ituarte, E. J., del Carmen, K., & Nedved, B. T. (2001). Metamorphic Competence, a Major Adaptive Convergence in Marine Invertebrate Larvae. American Zoologist, 41(5), 1123–1131. 10.1093/icb/41.5.1123

Hadfield, M., & Strathmann, M. (1996). Variability, flexibility and plasticity in life histories of marine invertebrates. 12.

Hao, Y., Hao, S., Andersen-Nissen, E., Mauck, W. M., Zheng, S., Butler, A., Lee, M. J., Wilk, A. J., Darby, C., Zager, M., Hoffman, P., Stoeckius, M., Papalexi, E., Mimitou, E. P., Jain, J., Srivastava, A., Stuart, T., Fleming, L. M., Yeung, B., … Satija, R. (2021). Integrated analysis of multimodal single-cell data. Cell, 184(13), 3573–3587.e29. 10.1016/j.cell.2021.04.048

Healy, J., & McInnes, L. (2024). Uniform manifold approximation and projection. Nature Reviews Methods Primers, 4(1), 82. 10.1038/s43586-024-00363-x

Hildebrandt, F., Otto, E., Rensing, C., Nothwang, H. G., Vollmer, M., Adolphs, J., Hanusch, H., & Brandis, M. (1997). A novel gene encoding an SH3 domain protein is mutated in nephronophthisis type 1. Nature Genetics, 17(2), 149–153. 10.1038/ng1097-149

Huelsenbeck, J. P., & Ronquist, F. (2001). MRBAYES: Bayesian inference of phylogenetic trees. Bioinformatics, 17(8), 754–755. 10.1093/bioinformatics/17.8.754

Kaul-Strehlow, S., Urata, M., Minokawa, T., Stach, T., & Wanninger, A. (2015). Neurogenesis in directly and indirectly developing enteropneusts: Of nets and cords. Organisms Diversity & Evolution, 15(2), 405–422.

Kolberg, L., Raudvere, U., Kuzmin, I., Adler, P., Vilo, J., & Peterson, H. (2023). g:Profiler— Interoperable web service for functional enrichment analysis and gene identifier mapping (2023 update). Nucleic Acids Research, 51(W1), W207–W212. 10.1093/nar/gkad347

Korsunsky, I., Millard, N., Fan, J., Slowikowski, K., Zhang, F., Wei, K., Baglaenko, Y., Brenner, M., Loh, P., & Raychaudhuri, S. (2019). Fast, sensitive and accurate integration of single-cell data with Harmony. Nature Methods, 16(12), 1289–1296. 10.1038/s41592-019-0619-0

Koushika, S. P., Lisbin, M. J., & White, K. (1996). ELAV, a Drosophila neuron-specific protein, mediates the generation of an alternatively spliced neural protein isoform. Current Biology : CB, 6(12), 1634–1641. 10.1016/s0960-9822(02)70787-2

Kuehn, E., Clausen, D. S., Null, R. W., Metzger, B. M., Willis, A. D., & Özpolat, B. D. (2021). Segment number threshold determines juvenile onset of germline cluster expansion in *Platynereis dumerilii*. Journal of Experimental Zoology Part B: Molecular and Developmental Evolution, jez.b.23100. 10.1002/jez.b.23100

Lacalli, T. C. (2005). Protochordate body plan and the evolutionary role of larvae: Old controversies resolved? Canadian Journal of Zoology, 83(1), 216–224. 10.1139/z04-162

Lacalli, T. C., & Gilmour, T. H. J. (2002). Locomotory and feeding effectors of the tornaria larva of *Balanoglossus biminiensis*: Tornaria structure and feeding. Acta Zoologica, 82(2), 117–126. 10.1046/j.1463-6395.2001.00075.x

Link, O., Jahnel, S. M., Janicek, K., Kraus, J., Montenegro, J. D., Zimmerman, B., Wick, B., Cole, A. G., & Technau, U. (2025). Changes of cell-type diversity in the polyp-to-medusa metagenesis of the scyphozoan jellyfish Aurelia coerulea (formerly sp.1). 10.1101/2023.08.24.554571

Littleton, J. T., Stern, M., Schulze, K., Perin, M., & Bellen, H. J. (1993). Mutational analysis of *Drosophila* synaptotagmin demonstrates its essential role in Ca(2+)-activated neurotransmitter release. Cell, 74(6), 1125–1134. 10.1016/0092-8674(93)90733-7

Lowe, C. J., Terasaki, M., Wu, M., Freeman, R. M., Runft, L., Kwan, K., Haigo, S., Aronowicz, J., Lander, E., Gruber, C., Smith, M., Kirschner, M., & Gerhart, J. (2006). Dorsoventral Patterning in Hemichordates: Insights into Early Chordate Evolution. PLoS Biology, 4(9), e291. 10.1371/journal.pbio.0040291

Marra, A. N., Adeeb, B. D., Chambers, B. E., Drummond, B. E., Ulrich, M., Addiego, A., Springer, M., Poureetezadi, S. J., Chambers, J. M., Ronshaugen, M., & Wingert, R. A. (2019). Prostaglandin signaling regulates renal multiciliated cell specification and maturation. Proceedings of the National Academy of Sciences, 116(17), 8409–8418. 10.1073/pnas.1813492116

Martín-Zamora, F. M., Liang, Y., Guynes, K., Carrillo-Baltodano, A. M., Davies, B. E., Donnellan, R. D., Tan, Y., Moggioli, G., Seudre, O., Tran, M., Mortimer, K., Luscombe, N. M., Hejnol, A., Marlétaz, F., & Martín-Durán, J. M. (2023). Annelid functional genomics reveal the origins of bilaterian life cycles. Nature, 615(7950), 105–110. 10.1038/s41586-022-05636-7

Maslakova, S. A. (2010). Development to metamorphosis of the nemertean pilidium larva. Frontiers in Zoology, 7(1), 30. 10.1186/1742-9994-7-30

Massri, A. J., Greenstreet, L., Afanassiev, A., Berrio, A., Wray, G. A., Schiebinger, G., & McClay, D. R. (2021). Developmental single-cell transcriptomics in the *Lytechinus variegatus* sea urchin embryo. Development, 148(19), dev198614. 10.1242/dev.198614

Miyamoto, N., Nakajima, Y., Wada, H., & Saito, Y. (2010). Development of the nervous system in the acorn worm *Balanoglossus simodensis*: Insights into nervous system evolution: Development of hemichordate nervous system. Evolution & Development, 12(4), 416–424. 10.1111/j.1525-142X.2010.00428.x

Morgan, T. H. (1891). The growth and metamorphosis of Tornaria. Journal of Morphology, 5, 407–358.

Myngbay, A., Manarbek, L., Ludbrook, S., & Kunz, J. (2021). The Role of Collagen Triple Helix Repeat-Containing 1 Protein (CTHRC1) in Rheumatoid Arthritis. International Journal of Molecular Sciences, 22(5). 10.3390/ijms22052426

Nakajima, Y., Humphreys, T., Kaneko, H., & Tagawa, K. (2004). Development and Neural Organization of the Tornaria Larva of the Hawaiian Hemichordate, *Ptychodera flava*. Zoological Science, 21(1), 69–78. 10.2108/0289-0003(2004)21%255B69:DANOOT%255D2.0.CO;2

Nielsen, C. (1998). Origin and evolution of animal life cycles. Biological Reviews, 73(2), 125–155. 10.1111/j.1469-185X.1997.tb00027.x

Nielsen, C. (2005). Larval and adult brains. Evolution & Development, 7(5), 483–489. 10.1111/j.1525-142X.2005.05051.x

Nielsen, C., & Hay-Schmidt, A. (2007). Development of the enteropneust *Ptychodera flava*: Ciliary bands and nervous system. Journal of Morphology, 268(7), 551–570. 10.1002/jmor.10533

Paganos, P., Ullrich-Lüter, J., Almazán, A., Voronov, D., Carl, J., Zakrzewski, A.-C., Zemann, B., Rusciano, M. L., Sancerni, T., Schauer, M., Akar, O., Caccavale, F., Cocurullo, M., Benvenuto, G., Croce, J. C., Lüter, C., & Arnone, M. I. (2025). Single Nucleus Profiling Highlights the All-Brain Echinoderm Nervous System. 10.1101/2025.03.24.644250

Paganos, P., Voronov, D., Musser, J. M., Arendt, D., & Arnone, M. I. (2021). Single-cell RNA sequencing of the Strongylocentrotus purpuratus larva reveals the blueprint of major cell types and nervous system of a non-chordate deuterostome. eLife, 10, e70416. 10.7554/eLife.70416

Pardos, F., & Benito, J. (1988). Blood Vessels and Related Structures in the Gill Bars of *Glossobalanus minutus* (Enteropneusta). Acta Zoologica, 69(2), 87–94. 10.1111/j.1463-6395.1988.tb00905.x

Park, J.-O., Pan, J., Möhrlen, F., Schupp, M.-O., Johnsen, R., Baillie, D. L., Zapf, R., Moerman, D. G., & Hutter, H. (2010). Characterization of the astacin family of metalloproteases in C. elegans. BMC Developmental Biology, 10(1), 14. 10.1186/1471-213X-10-14

Patry, W. L., Bubel, M., Hansen, C., & Knowles, T. (2020). Diffusion tubes: A method for the mass culture of ctenophores and other pelagic marine invertebrates. PeerJ, 8, e8938. 10.7717/peerj.8938

Perillo, M., Oulhen, N., Foster, S., Spurrell, M., Calestani, C., & Wessel, G. (2020). Regulation of dynamic pigment cell states at single-cell resolution. eLife, 9, e60388. 10.7554/eLife.60388

Peterson, K. J., Cameron, R. A., & Davidson, E. H. (1997). Set-aside cells in maximal indirect development: Evolutionary and developmental significance. BioEssays, 19(7), 623–631. 10.1002/bies.950190713

Pfeffer, P. L., Gerster, T., Lun, K., Brand, M., & Busslinger, M. (1998). Characterization of three novel members of the zebrafish Pax2/5/8 family: Dependency of Pax5 and Pax8 expression on the Pax2.1 (noi) function. Development, 125, 3063–3074.

Pham, K., & Hobert, O. (2019). Unlike Drosophila elav, the C. elegans elav orthologue exc-7 is not panneuronally expressed. microPublication Biology, 2019. 10.17912/micropub.biology.000189

Piovani, L., Leite, D. J., Guerra, L. A. Y., Simpson, F., Musser, J. M., Salvador-Martínez, I., Marlétaz, F., Jékely, G., & Telford, M. J. (2023). Single-cell atlases of two lophotrochozoan larvae highlight their complex evolutionary histories. Science Advances.

Robinow, S., & White, K. (1991). Characterization and spatial distribution of the ELAV protein during Drosophila melanogaster development. Journal of Neurobiology, 22(5), 443–461. 10.1002/neu.480220503

Ruppert, E. E., Cameron, C. B., & Frick, J. E. (1999). Endostyle-like Features of the Dorsal Epibranchial Ridge of an Enteropneust and the Hypothesis of Dorsal-Ventral Axis Inversion in Chordates. Invertebrate Biology, 118(2), 202–212. JSTOR. 10.2307/3227061

Rychel, A. L., Smith, S. E., Shimamoto, H. T., & Swalla, B. J. (2006). Evolution and development of the chordates: Collagen and pharyngeal cartilage. Molecular Biology and Evolution, 23(3), 541–549.

Samuelson, L. C., Wiebauer, K., Gumucio, D. L., & Meisler, M. H. (1988). Expression of the human amylase genes: Recent origin of a salivary amylase promoter from an actin pseudogene. Nucleic Acids Research, 16(17), 8261–8276. PubMed. 10.1093/nar/16.17.8261

Sánchez Alvarado, A., & Yamanaka, S. (2014). Rethinking Differentiation: Stem Cells, Regeneration, and Plasticity. Cell, 157(1), 110–119. 10.1016/j.cell.2014.02.041

Satija, R., Farrell, J. A., Gennert, D., Schier, A. F., & Regev, A. (2015). Spatial reconstruction of single-cell gene expression data. Nature Biotechnology, 33(5), 495–502. 10.1038/nbt.3192

Sebé-Pedrós, A., Chomsky, E., Pang, K., Lara-Astiaso, D., Gaiti, F., Mukamel, Z., Amit, I., Hejnol, A., Degnan, B. M., & Tanay, A. (2018). Early metazoan cell type diversity and the evolution of multicellular gene regulation. Nature Ecology & Evolution. 10.1038/s41559-018-0575-6

Sebé-Pedrós, A., Saudemont, B., Chomsky, E., Plessier, F., Mailhé, M.-P., Renno, J., Loe-Mie, Y., Lifshitz, A., Mukamel, Z., Schmutz, S., Novault, S., Steinmetz, P. R. H., Spitz, F., Tanay, A., & Marlow, H. (2018). Cnidarian Cell Type Diversity and Regulation Revealed by Whole-Organism Single-Cell RNA-Seq. Cell, 173(6), 1520–1534.e20. 10.1016/j.cell.2018.05.019

Simakov, O., Kawashima, T., Marlétaz, F., Jenkins, J., Koyanagi, R., Mitros, T., Hisata, K., Bredeson, J., Shoguchi, E., Gyoja, F., Yue, J. X., Chen, Y. C., Freeman, R. M., Sasaki, A., Hikosaka-Katayama, T., Sato, A., Fujie, M., Baughman, K. W., Levine, J., … Gerhart, J. (2015). Hemichordate genomes and deuterostome origins. Nature, 527(7579), 459–465. 10.1038/nature16150

Sogabe, S., Hatleberg, W. L., Kocot, K. M., Say, T. E., Stoupin, D., Roper, K. E., Fernandez-Valverde, S. L., Degnan, S. M., & Degnan, B. M. (2019). Pluripotency and the origin of animal multicellularity. Nature, 570(7762), 519–522. 10.1038/s41586-019-1290-4

Strathmann, R., & Bonar, D. (1976). Ciliary feeding of tornaria larvae of *Ptychodera flava* (Hemichordata: Enteropneusta). Marine Biology, 34(4), 317–324. 10.1007/BF00398125

Strathmann, R. R. (2020). Multiple origins of feeding head larvae by the Early Cambrian. Canadian Journal of Zoology, 98(12), 761–776. 10.1139/cjz-2019-0284

Stuart, T., Butler, A., Hoffman, P., Hafemeister, C., Papalexi, E., Mauck, W. M., Hao, Y., Stoeckius, M., Smibert, P., & Satija, R. (2019). Comprehensive Integration of Single-Cell Data. Cell, 177(7), 1888–1902.e21. 10.1016/j.cell.2019.05.031

Tagawa, K., Humphreys, T., & Satoh, N. (1998). Novel pattern of Brachyury gene expression in hemichordate embryos. Mechanisms of Development, 75(1–2), 139–143. 10.1016/S0925-4773(98)00078-1

Tarashansky, A. J., Xue, Y., Li, P., Quake, S. R., & Wang, B. (2019). Self-assembling manifolds in single-cell RNA sequencing data. eLife, 8, e48994. 10.7554/eLife.48994

Taylor, P. R., Martinez-Pomares, L., Stacey, M., Lin, H.-H., Brown, G. D., & Gordon, S. (2005). Macrophage receptors and immune recognition. Annual Review of Immunology, 23, 901–944. 10.1146/annurev.immunol.23.021704.115816

Truman, J. W., Price, J., Miyares, R. L., & Lee, T. (2023). Metamorphosis of memory circuits in Drosophila reveals a strategy for evolving a larval brain. eLife, 12, e80594. 10.7554/eLife.80594

Vasta, G. R., Amzel, L. M., Bianchet, M. A., Cammarata, M., Feng, C., & Saito, K. (2017). F-Type Lectins: A Highly Diversified Family of Fucose-Binding Proteins with a Unique Sequence Motif and Structural Fold, Involved in Self/Non-Self-Recognition. Frontiers in Immunology, 8, 1648–1648. PubMed. 10.3389/fimmu.2017.01648

Wray, G. A. (2000). The evolution of embryonic patterning mechanisms in animals. Seminars in Cell and Developmental Biology, 11(6), 385–393. 10.1006/scdb.2000.0191

Yoder, B. K., Hou, X., & Guay-Woodford, L. M. (2002). The Polycystic Kidney Disease Proteins, Polycystin-1, Polycystin-2, Polaris, and Cystin, Are Co-Localized in Renal Cilia. Journal of the American Society of Nephrology, 13(10), 2508. 10.1097/01.ASN.0000029587.47950.25

Zhao, J., Wang, G., Ming, J., Lin, Z., Wang, Y., The Tabula Microcebus Consortium, Agarwal, S., Agrawal, A., Al-Moujahed, A., Alam, A., Albertelli, M. A., Allegakoen, P., Ambrosi, T., Antony, J., Artandi, S., Aujard, F., Awayan, K., Baghel, A., Bakerman, I., … Yang, C. (2022). Adversarial domain translation networks for integrating large-scale atlas-level single-cell datasets. Nature Computational Science, 2(5), 317–330. 10.1038/s43588-022-00251-y

Zheng, G. X. Y., Terry, J. M., Belgrader, P., Ryvkin, P., Bent, Z. W., Wilson, R., Ziraldo, S. B., Wheeler, T. D., McDermott, G. P., Zhu, J., Gregory, M. T., Shuga, J., Montesclaros, L., Underwood, J. G., Masquelier, D. A., Nishimura, S. Y., Schnall-Levin, M., Wyatt, P. W., Hindson, C. M., … Bielas, J. H. (2017). Massively parallel digital transcriptional profiling of single cells. Nature Communications, 8(1), 14049. 10.1038/ncomms14049

Zheng, Y., Blair, D., & Bradley, J. E. (2013). Phyletic Distribution of Fatty Acid-Binding Protein Genes. PLoS ONE, 8(10), e77636. 10.1371/journal.pone.0077636

Zibetti, C., Adamo, A., Binda, C., Forneris, F., Toffolo, E., Verpelli, C., Ginelli, E., Mattevi, A., Sala, C., & Battaglioli, E. (2010). Alternative splicing of the histone demethylase LSD1/KDM1 contributes to the modulation of neurite morphogenesis in the mammalian nervous system. The Journal of Neuroscience : The Official Journal of the Society for Neuroscience, 30(7), 2521–2532. 10.1523/JNEUROSCI.5500-09.2010

